# EnZight: A Structure-Guided Algorithm to Identify and Prioritize Substitution Hotspots for Enzyme Engineering

**DOI:** 10.64898/2026.09.10.750651

**Authors:** Rune Rahbek Østergaard, Mikkel Lyskjær Jensen, Suzana Siebenhaar, Deniz Bicer, Peter Wad Sackett, Alexander Andersen, Matteo Tiberti, Elena Papaleo, Serina Robinson, Søren Skou Thirup, Peter Westh, Laura Rotilio, Jens Preben Morth

**Affiliations:** EnZync Center for Enzymatic Deconstruction of Thermoset Plastics; Biotechnology and Biomedicine, Technical University of Denmark, Søltofts Plads, DK-2800, Kongens Lyngby, Denmark; Interdisciplinary Nanoscience Center (iNANO), Aarhus University, Gustav Wields Vej 14, DK-8000, Aarhus, Denmark; Department of Health Technology, Technical University of Denmark, Ørsteds Plads, DK-2800, Kongens Lyngby, Denmark; Cancer Structural Biology, Danish Cancer Institute, Strandboule-varden 49, DK-2100 Copenhagen, Denmark; Institute of Aquatic Science and Technology CH-8600 Dübendorf, Switzerland; Institute for Biogeochemistry and Pollutant Dynamics, ETH Zürich, Universitätsstrasse 16, CH-8092 Zürich, Switzerland

**Keywords:** Enzyme engineering, Plastic degradation, Amidase, Structural bioinformatics

## Abstract

Homologous protein structures contain valuable information about tolerated sequence variation. However, translating this information into practical enzyme design strategies remains challenging. Here we present EnZight, a user-friendly web server that integrates homologous structural alignment with intuitive visualization to identify substitution hotspots in protein cores. EnZight exploits structurally aligned homologs to identify positions where the surrounding structural environment is conserved while the residue at the position varies across homologs. This enables prediction of substitutions that preserve fold integrity while modulating function and thermostability. The approach further provides interactive structural outputs that allow users to inspect and prioritize substitutions manually. To validate the use of EnZight, we used a polyurethane-degrading amidase as a proof of concept. We constructed 34 variants, and 97 % were successfully expressed, indicating high foldability of the predicted substitutions. Several substitutions improved both catalytic turnover and thermostability, and, importantly, beneficial substitutions combined additively, enabling stepwise accumulation of improvements. The best triple mutant variant exhibited a six-fold increase in catalytic turnover and 2 ^*°*^C increase in apparent melting temperature. Enhanced activity toward the pharmaceutical micropollutant flutamide further demonstrates EnZight’s broad applicability in identifying substitutions that enable enzyme optimization across diverse substrates.

## Introduction

Enzymes are essential for a broad range of biotechnology applications, such as additives in washing powder for lipid removal and biological waste treatment ^[1]^ biocatalysis of polysaccharides for biofuel ^[2]^, and enzymatic plastic recycling ^[3,4]^. However, the wild-type enzymes that embed the initial desired activity often lack the stability and promiscuous activity needed for efficient usage ^[3–7]^. Improving enzymatic stability and activity is therefore a major goal in enzyme engineering, as the endeavor will include designing enzymes for unnatural or industrial substrates ^[8]^. Depending on the application, enzymes may need to be efficient at high temperatures, under alkaline/acidic conditions, or in the presence of organic solvents ^[3–7]^. This is exemplified by plastic degrading enzymes, which have shown great improvements when optimized to tolerate high temperatures ^[4]^ or in the presence of organic solvents ^[3]^.

Evolutionary information has emerged as a powerful guide for protein engineering^[9,10]^. Since enzymes have been shaped by evolutionary selection over millions of years, conserved sequence positions often indicate residues that are important for maintaining protein structure, catalytic function, or substrate recognition. Computational tools such as ConSurf use sequence-based multiple sequence alignments (MSAs) to map sequence conservation onto protein structures, thereby helping to identify residues relevant for protein engineering ^[10]^.

Evolutionary sequence-based information is also widely used as an input for computational protein design. Stabilization tools such as PROSS and FireProt combine evolutionary information with physics-based energy calculations to identify mutations that are predicted to improve stability while minimizing disruption to protein structure and function. These approaches rely on the accumulation of multiple, spatially distributed mutations with additive stabilizing effects^[11,12]^. Other tools, such as FuncLib, use a similar approach for active-site engineering by generating focused libraries designed to alter catalytic properties ^[13]^. With the emergence of AlphaFold, ESMfold, and similar 3D prediction programs, the number of accurate protein structure models has expanded dramatically ^[14–19]^. This has created new opportunities to use structural information directly in protein engineering workflows. Some of which have already been combined with the enzyme engineering efforts. HotSpot Wizard integrates sequence, structural, and energy-based analysis to identify mutagenesis hotspots for engineering enzyme activity, specificity, and stability ^[20]^. Inverse-folding models, including ProteinMPNN and ESM-IF, have enabled the generation of protein sequences compatible with a given backbone structure ^[21–23]^. While these methods can produce highly stable proteins, preserving native activity is often challenging ^[24,25]^. However, recent work has shown that combining inverse folding with evolutionary information can help overcome this limitation, as exemplified by engineered TEV protease variants with improved stability, activity, and solubility ^[26]^.

Selecting a small number of substitutions that improve both enzyme stability and activity remains a challenge. Large mutant libraries can help identify beneficial variants, but they are often impractical when screening capacity is limited ^[27]^. Moreover, many substitutions disrupt structural integrity and result in loss-of-function variants, reducing the proportion of useful candidates in libraries. Prioritizing substitutions that are less likely to disrupt protein structure may, therefore, enable the design of smaller, more focused libraries enriched in functional variants.

Here, we introduce EnZight, a tool for generating structural alignments and structure-based multiple sequence alignments (SB-MSAs) to assess conservation across homologous proteins. By combining structural and conservation information, EnZight identifies amino acid substitutions within the protein core that are less likely to disrupt protein structure and can be used to design smaller and more focused mutant libraries. In addition, EnZight offers a user-friendly web server that makes analysis straightforward to run and resulting outputs easy to inspect.

We apply EnZight to a polyurethane-degrading signature amidase and demonstrate that it can accurately identify structurally tolerated substitutions that alter thermostability and catalytic turnover. The beneficial effects of the tested substitutions we show are additive. This allows stability and activity improvements to be accumulated stepwise without compromising structural integrity. Structural characterization indicated that the overall fold was preserved, while enhanced catalytic performance was maintained across distinct substrates, supporting the robustness and practical utility of the approach implemented through the use of EnZight.

## Results and Discussion

### EnZight is a Structure-Based Alignment and Residue Substitution Prediction Tool for Proteins

From a query structure, which can be either a computationally predicted model or an experimental model, En-Zight (https://services.healthtech.dtu.dk/services/EnZight-1.0/) predicts substitutions with a low risk of disrupting protein structure; we will refer to these as substitution hotspots. These include the candidate substitution position and the direct environment around the side chain in question. EnZight achieves this by structurally aligning the query structure with multiple homologous structural models, identified through Foldseek ^[28]^ or provided by the user (Figure 1a). Based on the structural alignment, a conservation score (see Methods) is computed for each residue, ranging from 0 for a completely variable position to 100 for a completely conserved position. In addition to generating conservation scores, EnZight produces a structure-based multiple sequence alignment (SB-MSA) by refining a conventional sequence-based MSA with gaps derived from the structural alignment. This refinement is especially beneficial in regions with low sequence similarity that are still structurally conserved.

**Figure 1.**
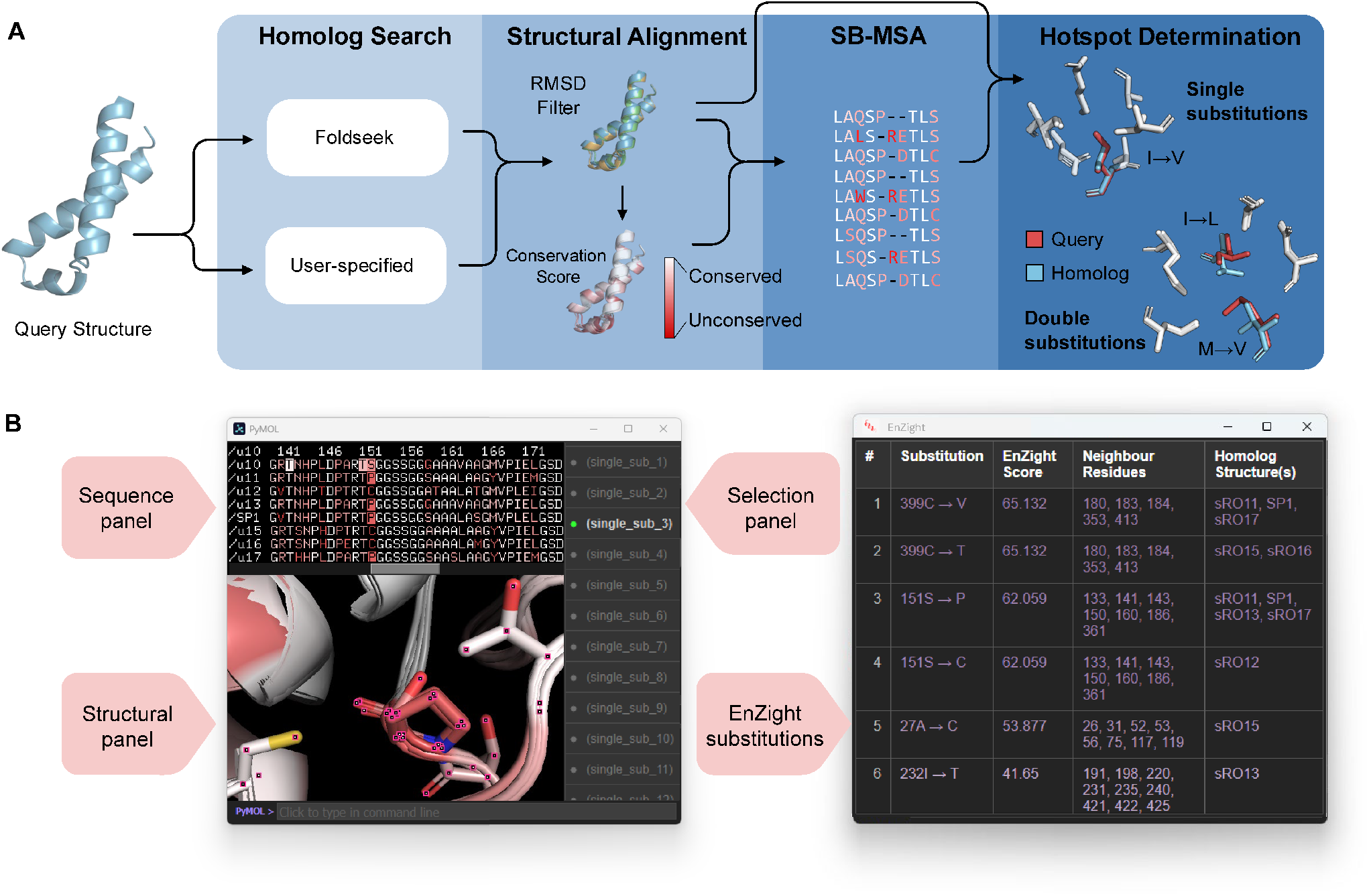
Program overview and example outputs of EnZight. a) A user-provided query structure is submitted to the homolog search module, which retrieves homologous structures via Foldseek ^[28]^ or accepts user-specified structures. Each homolog is structurally aligned to the query, and poorly aligned models are removed based on an RMSD threshold (default: 5 Å). From the resulting alignments, EnZight computes per-residue conservation score, then color-codes and computes a structural-based multiple sequence alignment (SB-MSA) using structural context to a sequence-based MSA. Finally, structural and sequence features are applied in the hotspot determination module to scan the protein core for positions where the query and a homolog differ in amino acid identity but share an equivalent local environment. Such positions are flagged as substitution hotspots, and the method identifies single- and double-substitution hotspots. b) Example output that can be visualized in PyMOL ^[29]^ separated into three panels: Sequence, structural, and selection panel. EnZight substitutions are shown in a tabular format, sorted and color-coded based on EnZight score.

The substitution hotspots are determined from the SBMSA and the structural alignment. Each substitution hotspot defines a position that differs between the query and at least one homolog while retaining an equivalent local environment. We hypothesized that equivalent local environments imply similar cavity geometry, so the substitution identified from the SB-MSA is likely to preserve the overall fold and structural integrity. The substitution hotspots are intended to focus primarily on the protein core, where predictions of the local structural environment are generally reliable ^[30]^. Thus, enabling the use of predicted structures as homologous inputs for substitution hotspot identification. EnZight also supports the identification of coupled double substitutions, defined as pairs of variable residues whose combined structural neighborhood is equivalent to that of at least one homolog.

EnZight generates two complementary outputs (Figure 1b). The first output is an interactive, PyMOL^[29]^-readable visualization that enables intuitive exploration of sequence–structure relationships. PyMOL is organized into three synchronized panels: a sequence panel, a structure panel, and a selection panel. Selecting a substitution hotspot in the selection panel automatically highlights the corresponding position in both the sequence and three-dimensional structure panels, enabling rapid inspection of the local structural environment for EnZight substitution candidates.

The second output is a table of allowed substitutions. Each entry represents either a single- or double-point substitution and is ranked according to its EnZight score (see Methods). This score reflects the combined conservation of the substituted position and its local structural environment, with higher scores indicating greater variability. Substitutions with high EnZight scores are therefore prioritized as potentially more structurally tolerated.

Taken together, these outputs provide an intuitive, interactive interface for structural analysis and a ranked list of substitution hotspots that can be used directly to guide experimental mutagenesis. Since EnZight uses existing structural predictions and requires relatively little computation, results are typically obtained within a few minutes. In addition, the user-friendly web interface makes EnZight easily accessible to users without extensive computational expertise.

### Experimental Validation of EnZight Yields Improved Substitutions with Additive Effects

To assess the quality of the substitution hotspot predictions from EnZight, we applied it to the amidase u10, an enzyme that is known to cleave urethane bonds in polyurethane plastics ^[31]^. EnZight identified 36 single-site substitution hotspots in u10 (Table S1) with EnZight conservation scores ranging from 3 to 66. To test whether EnZight scores correlate with experimental effects of substitutions, we attempted to generate constructs with all the predicted substitution hotspots. Of these, 34/36 (94 %) variants were successfully cloned, and 33/34 (97 %) were successfully expressed in *Escherichia coli* BL21 (DE3). The high expression success indicates that EnZight-guided substitutions likely had little effect on the folding and protein solubility, consistent with the intended minimal disruption of the overall protein architecture.

Enzymatic performance of the variants was measured by 2-hour kinetic endpoint measurements, and thermal stability was assessed by Nano Differential Scanning Fluorimetry (NanoDSF, Table S1). Hydrolysis according to the reaction in Figure 2a was used as a model for polyurethane degradation. For each mutant and the wild-type, the endpoint concentration of MUE–MDA (mM) and apparent melting temperature (*T*_*m,app*_, ^*°*^C) were plotted in a 2D plot (Figure 2b) to visualize the effect on both parameters. Except for three substitutions (M76I, V124L, and T410G), all expressed single-site variants retained measurable activity close to that of the wild type, and most displayed *T*_*m,app*_ values comparable to the wild type, with six outliers showing decreased stability. If, as hypothesized, these point substitutions do not perturb the global fold, combinations of spatially separated substitutions should have an additive effect on stability and activity. To test this, we constructed all pairwise and triple combinations of the four most favorable single substitutions – V44I, H90T, S151C, and S151P – and evaluated their thermal stability (*T*_*m,app*_) and endpoint activity ([MUE–MDA]) as described above (Figure 2b, Figure S1). Strikingly, for every combination, the observed change in Δ*T*_*m,app*_ was equal to the sum of the Δ*T*_*m,app*_ values of the corresponding individual substitutions, consistent with fully additive stabilization (Table 1). Additionally, all substitution combinations maintained apparent activity relative to wild-type in an additive manner. Collectively, these results demonstrate that EnZight-predicted hotspots likely can be combined additively to deliver simultaneous gains in both stability and activity.

**Table 1.** Additive effect of combinations of EnZight substitutions compared to wild-type. Experimental and predicted effect on *T*_*m,app*_ and MUE–MDA production for combinations of beneficial EnZight substitutions in u10. Values are reported as mean *±* SD. Predicted values are calculated as the sum of the effect on the single mutations contributing to a certain variant. Abbreviations: Mono-urethane ethylene methylenedianiline (MUE–MDA), predicted (Pred).

| Variant | $\Delta T_{m,app}$ (°C) | Pred $\Delta T_{m,app}$ (°C) | $\Delta$[MUE-MDA] (mM) | Pred $\Delta$[MUE-MDA] (mM) |
| --- | --- | --- | --- | --- |
| V44I | +1.01 ± 0.36 |  | -0.01 ± 0.05 |  |
| H90T | +0.80 ± 0.23 |  | +0.07 ± 0.05 |  |
| S151C | +0.89 ± 0.30 |  | 0.00 ± 0.05 |  |
| S151P | +0.19 ± 0.30 |  | +0.19 ± 0.05 |  |
| V44I_H90T | +1.69 ± 0.25 | +1.81 | +0.04 ± 0.08 | +0.06 |
| V44I_S151P | +1.42 ± 0.29 | +1.20 | +0.18 ± 0.05 | +0.18 |
| H90T_S151P | +1.26 ± 0.23 | +0.99 | +0.22 ± 0.05 | +0.26 |
| V44I_S151C | +2.03 ± 0.23 | +1.90 | -0.07 ± 0.05 | -0.01 |
| H90T_S151C | +1.89 ± 0.22 | +1.69 | +0.04 ± 0.05 | +0.07 |
| V44I_H90T_S151P | +2.00 ± 0.26 | +2.00 | +0.22 ± 0.05 | +0.25 |
| V44I_H90T_S151C | +2.52 ± 0.27 | +2.70 | +0.04 ± 0.05 | +0.06 |

**Figure 2.**
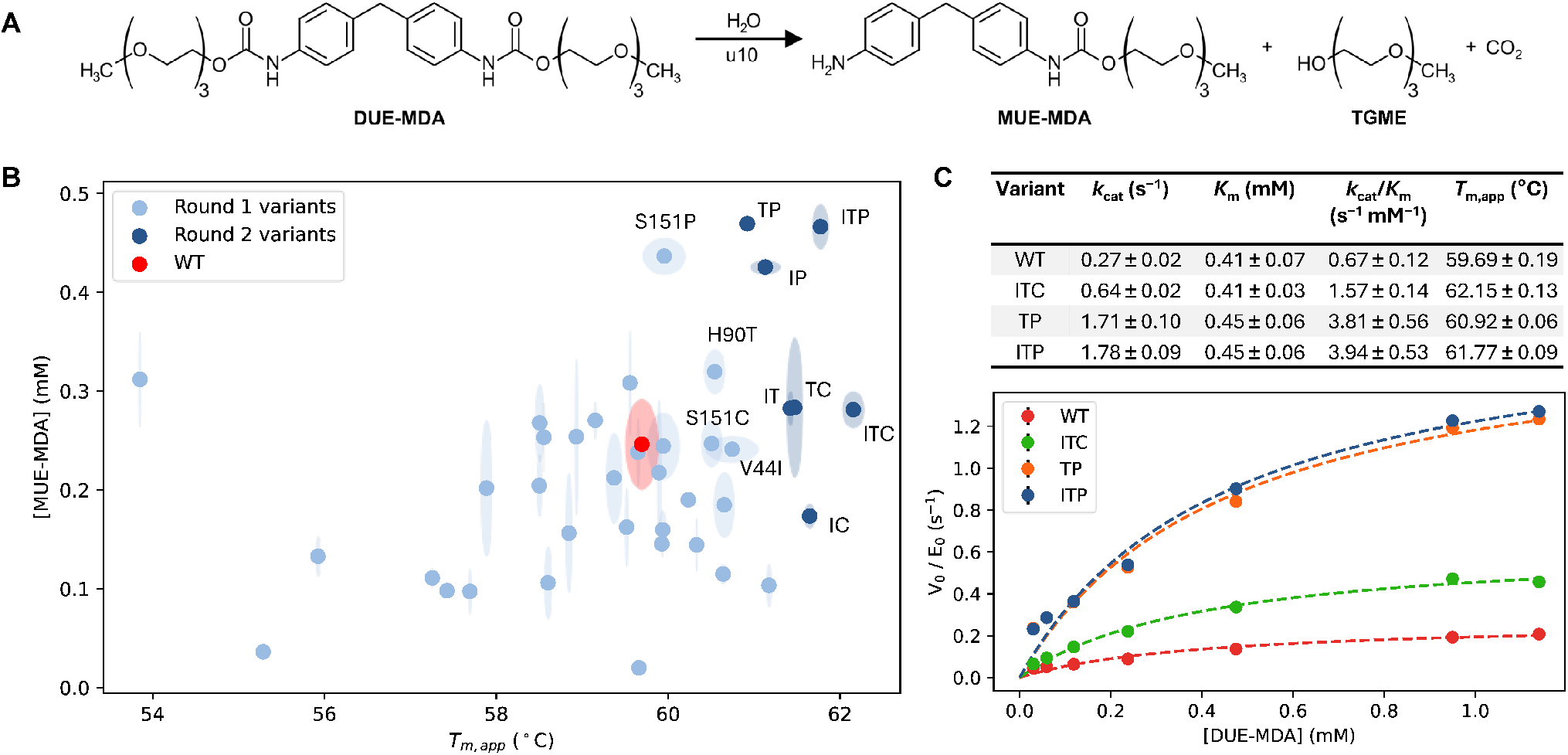
Stability and activity on DUE–MDA of u10 variants. a) The reaction of DUE–MDA degradation to MUE–MDA by u10. b-c) Wild-type u10 and engineered variants (round 1: 33 single-point substitutions; round 2: 7 combinatorial substitutions) were evaluated by b) a 2D plot of endpoint activity – quantified by MUE–MDA formation – versus apparent melting temperature (*T*_*m,app*_), where standard deviation (SD) is depicted as semi-transparent circles surrounding each data point; c) Michaelis–Menten kinetics for selected variants; SD shown as black error bars. Kinetic parameters reported as mean *±* SD. Abbreviations: Di-urethane ethylene methylenedianiline (DUE–MDA), mono-urethane ethylene methylenedianiline (MUE–MDA), triethylene glycol monomethyl ether (TGME), V44I_H90T (IT), V44I_S151C (IC), V44I_S151P (IP), H90T_S151C (TC), H90T_S151P (TP), V44I_H90T_S151C (ITC), V44I_H90T_S151P (ITP).

To obtain a more detailed understanding of the effects of substitutions, we performed Michaelis–Menten analysis on the wild-type and the most promising variants: H90T_S151P (TP), V44I_H90T_S151P (ITP), and V44I_H90T_S151C (ITC) (Figure 2c). The catalytic efficiency (*k*_*cat*_/*K*_*m*_) of TP and ITP were the highest with 3.81 *±* 0.56 s^−1^mM^−1^ and 3.94 *±* 0.53 s^−1^mM^−1^, respectively. This is *∼*6-fold higher than the wild-type *k*_*cat*_/*K*_*m*_ of 0.67 *±* 0.12 s^−1^mM^−1^. The ITC triple substitution had a *k*_*cat*_/*K*_*m*_ of 1.57 *±* 0.14 s^−1^mM^−1^ between the *k*_*cat*_/*K*_*m*_-values of the wild-type and the other substitutions, suggesting that both the H90T and the S151P have a positive effect on *k*_*cat*_/*K*_*m*_ for DUE–MDA degradation. Since TP and ITP had similar *k*_*cat*_/*K*_*m*_ values, it seems that V44I does not affect activity, which is consistent with the endpoint kinetic data (Figure 2b). It is noteworthy that *K*_*m*_ was unaffected for all the variants, suggesting that the substitutions in the most promising variants stabilize the transition state for the chemical step (carbamate bond cleavage) while leaving the free energy of the ES ground state unchanged.

### Screening Showed Substrate-independent Effects of Substitutions

The foundation of our approach is to preserve and optimize the structural integrity of the fold by targeting specific hotspot regions; our approach should therefore also be largely substrate-independent. To address this question, we assessed whether the increased catalytic efficiency (*k*_*cat*_/*K*_*m*_) was specific to the polyurethane model substrate DUE–MDA, or if similar kinetic parameters could be determined by using additional substrates. The wild-type and the most promising variants – H90T_S151P (TP), V44I_H90T_S151P (ITP), and V44I_H90T_S151C (ITC) – were evaluated using four *para*-nitrophenyl (*p*NP) substrates, *p*NP-butyrate, *p*NP-butanamide, and ethyl *p*NP-carbamate (Figure 3a-c). These substrates share similar molecular weight and overall structure but differ in the scissile bond type (ester, amide, and carbamate respectively).

**Figure 3.**
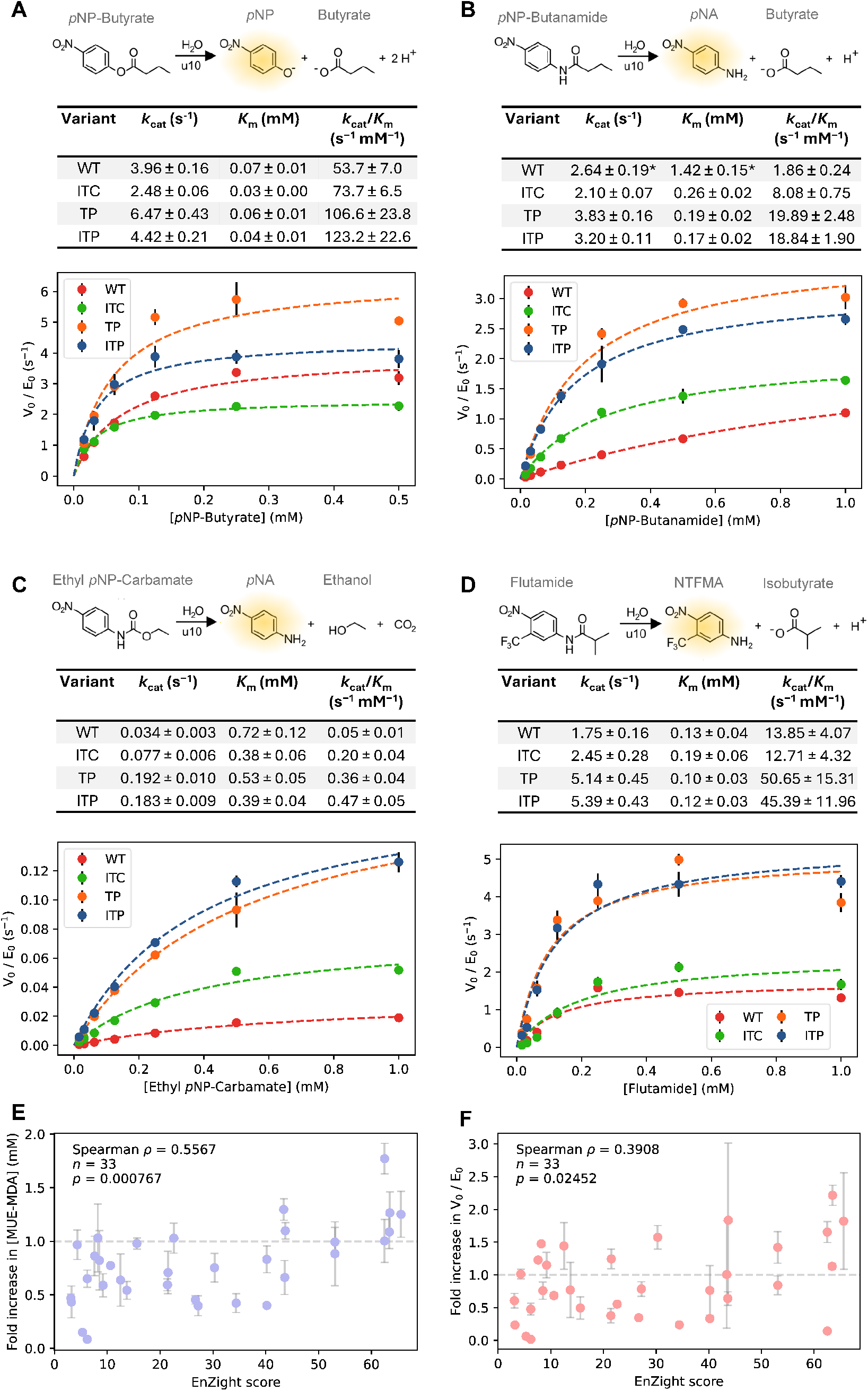
Substrate screening of u10 variants. Chemical reaction, kinetic parameters, and Michaelis–Menten plots using u10 variants on a) *p*NP-Butyrate, b) *p*NP-Butanamide, c) Ethyl *p*NP-Carbamate, and d) flutamide. e-f) Fold increase in activity as a function of EnZight score for all tested EnZight single-point substitutions of u10 on e) DUE–MDA and f) flutamide. All data points are reported as mean *±* SD from technical triplicates. Statistical parameters (Spearman *ρ*, number of samples (n), p-value) for the correlation were included. Horizontal line (*y* = 1) was included as wild-type activity. Abbreviations: *para*-nitrophenol (*p*NP), *para*-nitroaniline (*p*NA), *para*-nitro-*meta*-(trifluoromethyl)aniline (NTFMA). *Marks kinetic parameters that are not reliable due to almost linear correlation of initial rates.

Across all substrates catalytic efficiency decreased with increasing bond stability, as expected ^[32,33]^. The highest *k*_*cat*_/*K*_*m*_ values observed for the ester substrate (up to 123.2 *±* 22.6 s^−1^mM^−1^), followed by the amide (up to 19.89 *±* 2.48 s^−1^mM^−1^), and carbamate substrates (up to 0.47 *±* 0.05 s^−1^mM^−1^). For the ester substrate *p*NP-butyrate, only minor differences in catalytic performance were observed between the wild type enzyme and the variants, with the best-performing variant (ITP) showing a *∼*2.3-fold increase in *k*_*cat*_/*K*_*m*_ relative to wild-type (Figure 3a). In contrast, the Michaelis–Menten plot for the amide substrate *p*NP-butanamide (Figure 3b) shows a significantly larger difference between the wild-type and the best-performing variant (TP) with a *∼*10-fold increase in specificity constant (*k*_*cat*_/*K*_*m*_). Notably, the *k*_*cat*_ values obtained with ethyl *p*NP-carbamate follow the same trend as those measured for DUE–MDA (Figure 3c); however, they are uniformly *∼*10-fold lower in absolute value. As with DUE–MDA, the best-performing variants on ethyl *p*NP-carbamate (TP and ITP) showed a *∼*5.5-fold increase in *k*_*cat*_ relative to the wild-type u10. These results suggest that *p*NP-carbamate can serve as a colorimetric substrate to mimic DUE–MDA degradation, which also has the same scissile bond.

The observed activity towards ester, amide, and carbamate substrates indicates that u10 displays pronounced catalytic promiscuity. To explore whether this promiscuity extends to environmentally relevant small molecules, we examined the enzyme’s activity on pharmaceutical micropollutants containing amide functionalities. As a representative compound, the antiandrogen drug flutamide, a drug widely used in chemotherapy for prostate cancer and frequently detected as a pharmaceutical micropollutant in aquatic environments ^[34,35]^, was selected as a model substrate (Figure 3d). The ability to cleave such compounds highlights the potential of u10 as a candidate for enzymatic micropollutant removal.

All tested variants exhibited substantially higher catalytic efficiencies on flutamide than on the polyurethane model substrate DUE–MDA, with *k*_*cat*_/*K*_*m*_ values approximately 8–20-fold higher. The TP and ITP variants showed the highest efficiencies (50.65 *±* 15.31 and 45.39 *±* 11.96 s^−1^mM^−1^, respectively), corresponding to an approximately 3.5-fold improvement relative to the wild-type enzyme (*k*_*cat*_/*K*_*m*_ = 13.85 *±* 4.07 s^−1^mM^−1^). These results indicate that the effects of EnZight-guided substitutions are not substrate-dependent. Catalytic performance can be enhanced across multiple substrates, including polyurethane model substrates and pharmaceutical micropollutants.

### EnZight Score Correlates with Improved Substitution Effects

To examine whether the functional impact of substitutions correlates with EnZight conservation, the fold change in activity for all tested variants was plotted against their EnZight score (Figure 3e–f). Activity was quantified using end-point kinetics for DUE–MDA (1 mM) and observed turnover rates (V_0_/E_0_) for flutamide (0.45 mM). Significant positive correlations were observed between the EnZight score and activity enhancement for both DUE–MDA (Spearman’s *ρ* = 0.56, *n* = 33, *p* = 0.0007; Figure 3e) and flutamide (Spearman’s *ρ* = 0.39, *n* = 33, *p* = 0.025; Figure 3f), with the stronger association observed for DUE–MDA. Furthermore, the two positions associated with the greatest activity improvements for both substrates ranked among the highest-scoring positions in the EnZight output (Table S1). Taken together, these results indicate that substitutions at less conserved positions in a less conserved environment, corresponding to high EnZight scores, have a greater tendency to improve catalytic activity and that this trend is largely maintained across chemically distinct substrates. This enables substitutions with high EnZight scores to be prioritized, thereby allowing the design of smaller, more focused EnZight-guided libraries. No correlation between EnZight score and effect on *T*_*m,app*_ was observed (Figure S3).

### Structural Analysis Shows EnZight-guided Substitutions Affect Catalysis Through Local Interactions

The fundamental principle of EnZight-guided hotspot identification is that substitutions should minimally disrupt the overall protein structure and instead affect stability or catalytic activity through local interactions. This limited structural disruption is consistent with the additive improvements observed when favorable substitutions were combined. To investigate the structural basis of these increases in stability and catalytic activity, we determined the crystal structures of the wild-type enzyme and the ITP variant by X-ray crystallography. To stabilize the enzymes and prevent active-site disorder – a common issue in similar amidase-signature proteins ^[36]^ – a suicide inhibitor (LUI; Figure S4), recently described ^[37]^ and tested ^[38]^, was added prior to crystallization. The structure of the ITP variant in complex with LUI was determined at 2.2 Å resolution (Figure 4, Table S2, and Figure S5c), with the inhibitor covalently bound to the nucleophilic S178 of the K79-*cis*-S154-S178 catalytic triad (Figure S6). However, despite extensive screening, no crystals were obtained for the wild-type:LUI complex. Instead, we managed to determine the structure of the wild-type enzyme in the unbound state at a resolution at 1.8 Å (Figure S7a, Table S2, Figure S5a-b), although the active site was largely disordered. Consequently, of the three residues that comprise the triple substitution in ITP (positions 44, 90, and 151), only residue 44 was resolved in the wild-type structure, allowing experimental data comparison with the ITP structure only at this position, while other comparisons are performed using an Alphafold3 model of the wild type. Despite these limitations, the wild-type and the ITP variant showed high structural similarity, with an RMSD at 0.32 Å based on the C-alpha positions (Figure S7b).

**Figure 4.**
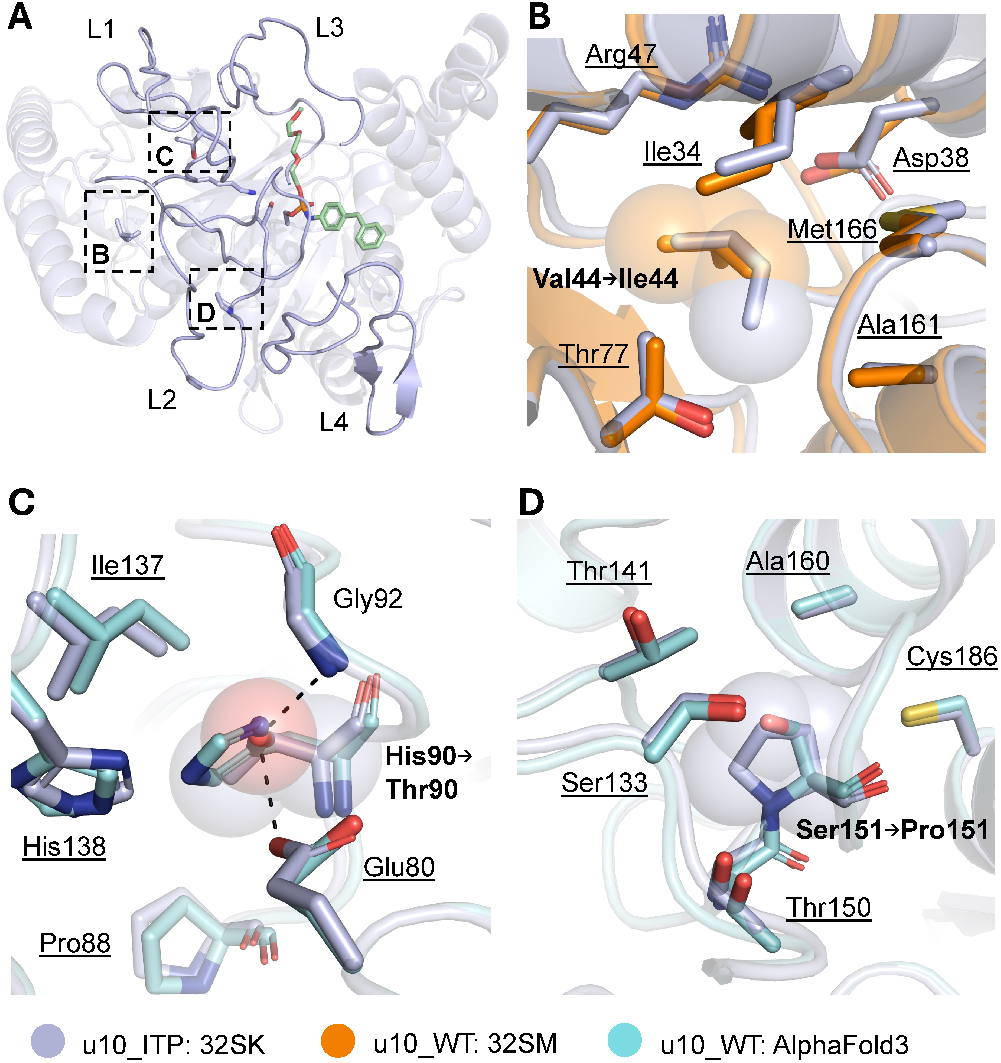
Structural analysis of the u10 ITP variant. Hydrogen bonds are shown as dashed lines (2.9 Å to 3.0 Å). a) Overall structure of the u10 ITP variant with the amidase signature loops highlighted (L1–L4). The LUI inhibitor is shown in green, and the side chains of the catalytic triad and the mutated residues are displayed as sticks. The location of the magnified views are indicated by dashed rectangles. b-d) Magnified views fo the local environment of b) the V44I substitution, c) the H90T substitution, and d) the S151P substitution. Residue labels indicate the corresponding substitution in bold, whereas neighboring residues predicted by EnZight are underlined. Residues shown without bold or underlined formatting were not predicted by EnZight to have side chains within the local environment of the substituted residue.

No displacement of neighboring residues was observed for any of the substitutions comprising the ITP triple variant (Figure 4b-d). This is consistent with the hypothesis that EnZight-guided substitutions are structurally tolerated and minimally disruptive. The V44I substitution fills a hydrophobic pocket through the introduction of an additional methyl group (Figure 4b), consistent with the observed increase in stability without compromising catalytic turnover. The H90T substitution, located in loop L1, introduces a stable hydrogen-bonding network that stabilizes E80 (Figure 4c). Since E80 is adjacent to the catalytic residue K79, this interaction may stabilize the local environment of the catalytic triad. Improved positioning of K79 could facilitate proton transfer and thereby enhance catalytic turnover. Finally, the S151P substitution is located in a small hydrophobic pocket (Figure 4d). This substitution may improve local hydrophobic packing and thereby stabilize the segment of loop L2 spanning residues 151–160, which contains *cis*-S154. Because *cis*-S154 plays a key role in the formation of the rate-limiting intermediate ^[39]^, improved positioning of this residue could lower the activation barrier of the catalytic step and thereby increase turnover.

This structural analysis shows how EnZight guided substitutions alter catalytic activity and thermostability through local interactions rather than through global structural perturbations. This is consistent with the high structural integrity of the variants and supports the premise of additive substitutions with minimal effect on the global protein architecture.

## Conclusion

It is widely accepted that residue conservation is a key parameter for protein engineering ^[9–13,40–42]^. Here, we present EnZight, a structural alignment tool that integrates sequence and structural information from homologous proteins to map residue conservation and enable intuitive visualization in PyMOL ^[29]^. By combining this conservation mapping with local structural context, EnZight identifies substitution hotspots that can be used for rational enzyme engineering. Using the polyurethane-degrading enzyme u10 ^[31]^ as a model system, we demonstrate that EnZight-guided substitutions can be used for the optimization of enzymatic activity and stability.

In the case of u10, favorable substitutions identified by EnZight displayed additive effects on both activity and stability when combined. The identified hotspots acted largely independently and did not exhibit epistatic interactions, a conclusion supported by our structural analysis. In many protein engineering strategies, substitutions interact in unpredictable ways, often resulting in non-additive or even deleterious effects when combined ^[43,44]^. In contrast, the additive behavior observed here enables a predictable optimization strategy in which improvements from individual substitutions can be accumulated stepwise. Such modularity simplifies exploration of the combinatorial sequence space and allows efficient optimization using a limited number of targeted substitutions.

EnZight-guided substitutions of u10 primarily increased the turnover number (*k*_*cat*_), whereas *K*_*m*_ remained relatively unchanged, indicating that apparent substrate affinity was not substantially affected. Similar activity trends were observed for substrates sharing the same scissile bond, regardless of their overall molecular structure, supporting the conclusion that the improvements were linked primarily to bond chemistry rather than substrate specificity. This suggests that simple, inexpensive colorimetric model substrates can be used to screen EnZight-guided substitutions for activity improvements during the early stages of enzyme engineering. More relevant and chemically complex substrates can then be introduced at later stages for detailed validation. Thus, enzyme optimization may be performed using high-throughput colorimetric assays, provided that the model substrate contains the relevant scissile bond.

A key finding was the correlation between the local environmental conservation (EnZight scores) and the functional effects of substitutions on enzymatic activity. Substitutions at less conserved environments were more likely to enhance catalytic performance, supporting residue conservation as a rational criterion for prioritizing mutagenesis targets. In contrast, no clear correlation between EnZight scores and changes in protein stability was observed in the present dataset. This suggests that, for u10, the EnZight score was more predictive of activity effects than stability changes. However, this observation does not exclude the use of EnZight-guided substitutions for stability engineering, as several substitutions nevertheless produced improvements in stability.

In this work, we identify substitution hotspots exclusively from naturally occurring homologs. However, EnZight is not restricted to evolutionary datasets and can be directly applied to ensembles of designed protein structures, such as those generated by inverse folding ^[21–23]^, by simply substituting the homolog input set with designed structures. In contrast to natural homologs, computationally designed proteins often show high thermodynamic stability^[26,45,46]^. Design ensembles, therefore, represent datasets enriched for stability-compatible sequence features. Applying EnZight in this context could enable the identification of substitutions that are consistently tolerated across stable designs, providing a complementary strategy for the rational discovery of stabilizing substitutions. This capability could make EnZight a useful framework for integrating natural sequence diversity with computational protein design and for combining these approaches with complementary protein-engineering tools.

## Supporting Information

The authors have cited additional references within the Supporting Information (Ref. ^[47–57]^).

## Acknowledgements

EnZync is supported by Challenge grant NNF22OC0072891 from the Novo Nordisk Foundation. We are grateful to the rest of the EnZync consortium members for constructive and helpful feedback. The X-ray diffraction data were collected at the Biomax beamline operated by MAXIV storage ring (MAXIV, Lund, Sweden), proposal number 20250330 (EnZync: Structural Characterization of Thermoset Plastic Degrading Enzymes). We thank Tobias Krojer for the help during data collection. We thank Catalyze4X for assistance with crystallization and shipping and data collection, grant NNF24OC0088714. We thank Martin Bundgaard Johansen, Andreas Sommerfeldt, Alexander Sandahl, and Nikolaj Lilholm Villadsen (Danish Technological Institute, Denmark) for synthesizing DUE–MDA, MUE–MDA, and the LUI inhibitor. We thank Peter Rahbek Østergaard for valuable discussions and guidance, particularly in establishing an efficient protein purification pipeline. ChatGPT (OpenAI) was used to support code development and manuscript editing. All generated suggestions were critically reviewed and verified by the authors, who take full responsibility for the final content.

## Entry for the Table of Contents

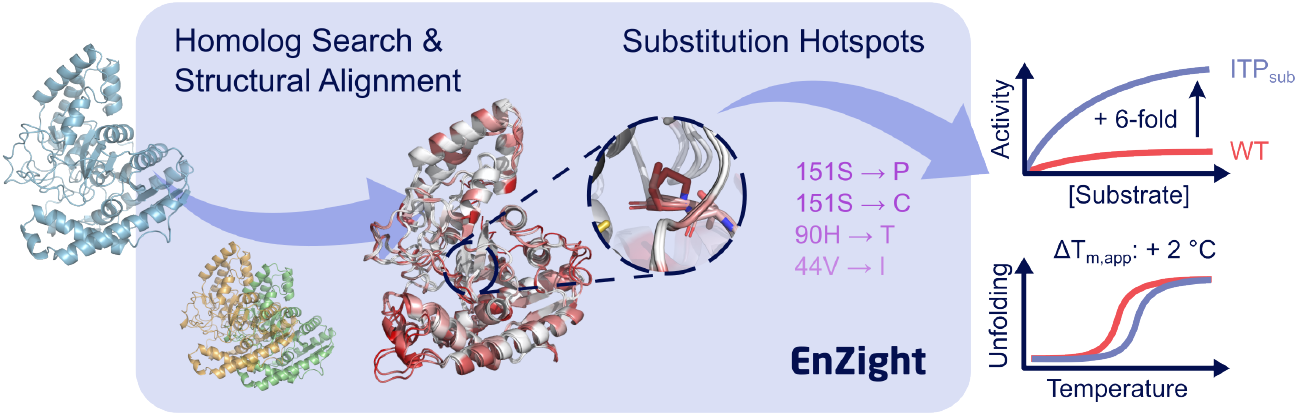

Homologous protein structures reveal amino acid substitutions that can be introduced without compromising structural integrity. EnZight structurally aligns homologs, visualizes residue conservation, and identifies substitution hotspots. Experimental validation showed that EnZight-guided substitutions improved thermostability and increased catalytic turnover up to six-fold, enabling rational, stepwise enzyme optimization.

## References

[1] A. Al-Ghanayem, B. Joseph, Applied Microbiology and Biotechnology 2020, 104, 2871.

[2] S. P. Gilmore, J. K. Henske, M. A. O’Malley, Bioengineered 2015, 6, 204.

[3] Y. Chen, J. Sun, K. Shi, T. Zhu, R. Li, R. Li, X. Liu, X. Xie, C. Ding, W.-C. Geng, et al., Science 2025, 390, 503.

[4] V. Tournier, C. M. Topham, A. Gilles, B. David, C. Folgoas, E. Moya-Leclair, E. Kamionka, M.-L. Desrousseaux, H. Texier, S. Gavalda, et al., Nature 2020, 580, 216.

[5] J. A. Littlechild, Frontiers in Bioengineering and Biotechnology 2015, 3, 161.

[6] K. S. Siddiqui, Critical Reviews in Biotechnology 2017, 37, 309.

[7] J. E. Gado, M. Knotts, A. Y. Shaw, D. Marks, N. P. Gauthier, C. Sander, G. T. Beckham, Nature Machine Intelligence 2025, 7, 716.

[8] Y. Huang, L. Wen, B. Yang, Journal of Agricultural and Food Chemistry 2026, 74, 185.

[9] W. P. Russ, M. Figliuzzi, C. Stocker, P. Barrat-Charlaix, M. Socolich, P. Kast, D. Hilvert, R. Monasson, S. Cocco, M. Weigt, et al., Science 2020, 369, 440.

[10] Yariv, E. Yariv, A. Kessel, G. Masrati, A. B. Chorin, E. Martz, I. Mayrose, T. Pupko, N. Ben-Tal, Protein Science 2023, 32, e4582.

[11] M. Musil, J. Stourac, J. Bendl, J. Brezovsky, Z. Prokop, J. Zendulka, T. Martinek, D. Bednar, J. Damborsky, Nucleic Acids Research 2017, 45, W393.

[12] A. Goldenzweig, M. Goldsmith, S. E. Hill, O. Gertman, P. Laurino, Y. Ashani, O. Dym, T. Unger, S. Albeck, J. Prilusky, et al., Molecular Cell 2016, 63, 337.

[13] O. Khersonsky, R. Lipsh, Z. Avizemer, Y. Ashani, M. Goldsmith, H. Leader, O. Dym, S. Rogotner, D. L. Trudeau, J. Prilusky, et al., Molecular Cell 2018, 72, 178.

[14] J. Jumper, R. Evans, A. Pritzel, T. Green, M. Figurnov, O. Ronneberger, K. Tunyasuvunakool, R. Bates, A. Žídek, A. Potapenko, et al., Nature 2021, 596, 583.

[15] J. Abramson, J. Adler, J. Dunger, R. Evans, T. Green, A. Pritzel, O. Ronneberger, L. Willmore, A. J. Ballard, J. Bambrick, et al., Nature 2024, 630, 493.

[16] Bertoni, M. Tsenkov, P. Magana, S. Nair, I. Pidruchna, M. Q. L. Afonso, A. Midlik, U. Paramval, D. Lawal, A. Tanweer, et al., Nucleic Acids Research 2026, 54, D358.

[17] B. Webb, A. Sali, Protein Structure Modeling with MODELLER, Springer, New York, NY 2017.

[18] Z. Lin, H. Akin, R. Rao, B. Hie, Z. Zhu, W. Lu, N. Smetanin, R. Verkuil, O. Kabeli, Y. Shmueli, et al., Science 2023, 379, 1123.

[19] M. Baek, F. DiMaio, I. Anishchenko, J. Dauparas, S. Ovchinnikov, G. R. Lee, J. Wang, Q. Cong, L. N. Kinch, R. D. Schaeffer, et al., Science 2021, 373, 871.

[20] L. Sumbalova, J. Stourac, T. Martinek, D. Bednar, J. Damborsky, Nucleic Acids Research 2018, 46, W356.

[21] J. Dauparas, I. Anishchenko, N. Bennett, H. Bai, R. J. Ragotte, L. F. Milles, B. I. M. Wicky, A. Courbet, R. J. de Haas, N. Bethel, et al., Science 2022, 378, 49.

[22] J. Dauparas, G. R. Lee, R. Pecoraro, L. An, I. Anishchenko, C. Glasscock, D. Baker, Nature Methods 2025, 22, 717.

[23] Hsu, R. Verkuil, J. Liu, Z. Lin, B. Hie, T. Sercu, A. Lerer, A. Rives, bioRxiv preprint (2022), DOI: 10.1101/2022.04.10.487779.

[24] Y. Liu, R. Wu, X. Wang, S. Wang, L. Chen, F. Li, Q. Chen, H. Liu, Nature Communications 2025, 16, 10177.

[25] P. Notin, N. Rollins, Y. Gal, C. Sander, D. Marks, Nature Biotechnology 2024, 42, 216.

[26] K. H. Sumida, R. Núñez-Franco, I. Kalvet, S. J. Pellock, B. I. M. Wicky, L. F. Milles, J. Dauparas, J. Wang, Y. Kipnis, N. Jameson, et al., Journal of the American Chemical Society 2024, 146, 2054.

[27] L. S. Vidal, M. Isalan, J. T. Heap, R. Ledesma-Amaro, RSC Chemical Biology 2023, 4, 271.

[28] M. van Kempen, S. S. Kim, C. Tumescheit, M. Mirdita, J. Lee, C. L. Gilchrist, J. Söding, M. Steinegger, Nature Biotechnology 2024, 42, 243.

[29] Schrödinger, LLC, The PyMOL Molecular Graphics System, Version 1.8 2015.

[30] K. T. Bæk, K. P. Kepp, Journal of Chemical Information and Modeling 2022, 62, 3391.

[31] L. Rotilio, R. R. Østergaard, E. M. Thiesen, P. Paiva, M. B. Johansen, A. Sommerfeldt, A. Sandahl, M. B. Keller, S. Siebenhaar, D. E. Otzen, et al., bioRxiv preprint (2026), DOI: 10.64898/2026.02.11.705263.

[32] P. Fugolin, M. G. Logan, A. J. Kendall, J. L. Ferracane, C. S. Pfeifer, Dental Materials 2021, 37, 805.

[33] T. M. Chapman, Journal of Polymer Science Part A: Polymer Chemistry 1989, 27, 1993.

[34] T. H. Miller, K. T. Ng, S. T. Bury, S. E. Bury, N. R. Bury, L. P. Barron, Environment International 2019, 129, 595.

[35] B. R. Goldspiel, D. R. Kohler, DICP 1990, 24, 616.

[36] T. Bayer, G. J. Palm, L. Berndt, H. Meinert, Y. Branson, L. Schmidt, C. Cziegler, I. Somvilla, C. Zurr, L. G. Graf, U. Janke, C. P. S. Badenhorst, S. König, M. Delcea, U. Garscha, R. Wei, M. Lammers, U. T. Bornscheuer, Angewandte Chemie International Edition 2024, 63, e202404492.

[37] L. Teixeira, P. Paiva, L. Rotilio, J. P. Morth, P. Fernandes, M. Ramos, chemRxiv preprint (2026), DOI: 10.26434/chemrxiv.15000233/v1.

[38] Bicer, D. Kochubei, R. Graham, S. Pena-Diaz, L. Rotilio, N. L. Villadsen, A. Sommerfeldt, M. B. Johansen, A. Sandahl, S. S. Thirup, J. P. Morth, D. E. Otzen, bioRxiv preprint (2026), DOI: 10.64898/2026.07.06.734427.

[39] P. Paiva, L. M. C. Teixeira, R. Wei, W. Liu, G. Weber, J. P. Morth, P. Westh, A. R. Petersen, M. B. Johansen, A. Sommerfeldt, et al., Chemical Science 2025, 16, 2437.

[40] D. Bednar, K. Beerens, E. Sebestova, J. Bendl, S. Khare, R. Chaloupkova, Z. Prokop, J. Brezovsky, D. Baker, J. Damborsky, PLoS Computational Biology 2015, 11, e1004556.

[41] J. J. Weinstein, A. Goldenzweig, S. Hoch, S. J. Fleishman, Bioinformatics 2021, 37, 123.

[42] H. Fei, Y. Li, Y. Liu, J. Wei, A. Chen, C. Gao, Cell 2025, 188, 4674.

[43] T. N. Starr, J. W. Thornton, Protein Science 2016, 25, 1204.

[44] R. Lipsh-Sokolik, S. J. Fleishman, Proceedings of the National Academy of Sciences 2024, 121, e2314999121.

[45] A.H.-W. Yeh, C. Norn, Y. Kipnis, D. Tischer, S. J. Pellock, D. Evans, P. Ma, G. R. Lee, J. Z. Zhang, I. Anishchenko, et al., Nature 2023, 614, 774.

[46] D. Baker, Protein Science 2019, 28, 678.

[47] T. U. Consortium, Nucleic Acids Research 2025, 53, D609.

[48] R. C. Edgar, Nature Communications 2022, 13, 6968.

[49] S. F. Altschul, T. L. Madden, A. A. Schäffer, J. Zhang, Z. Zhang, W. Miller, D. J. Lipman, Nucleic Acids Research 1997, 25, 3389.

[50] C. Camacho, G. Coulouris, V. Avagyan, N. Ma, J. Papadopoulos, K. Bealer, T. L. Madden, BMC Bioinformatics 2009, 10, 421.

[51] L. Rotilio, T. Bayer, H. Meinert, L. M. C. Teixeira, M. B. Johansen, A. Sommerfeldt, A. R. Petersen, A. Sandahl, M. B. Keller, J. Holck, et al., Angewandte Chemie International Edition 2025, 64, e202419535.

[52] W. Kabsch, Acta Crystallographica Section D: Biological Crystallography 2010, 66, 125.

[53] M.-F. Incardona, G. P. Bourenkov, K. Levik, R. A. Pieritz, A. N. Popov, O. Svensson, Journal of Synchrotron Radiation 2009, 16, 872.

[54] A. J. McCoy, R. W. Grosse-Kunstleve, P. D. Adams, M. D. Winn, L. C. Storoni, R. J. Read, Journal of Applied Crystallography 2007, 40, 658.

[55] P. Emsley, K. Cowtan, Acta Crystallographica Section D: Biological Crystallography 2004, 60, 2126.

[56] P. V. Afonine, R. W. Grosse-Kunstleve, N. Echols, J. J. Headd, N. W. Moriarty, M. Mustyakimov, T. C. Terwilliger, A. Urzhumtsev, P. H. Zwart, P. D. Adams, Acta Crystallographica Section D: Biological Crystallography 2012, 68, 352.

[57] V. B. Chen, W. B. Arendall, J. J. Headd, D. A. Keedy, R. M. Immormino, G. J. Kapral, L. W. Murray, J. S. Richardson, D. C. Richardson, Acta Crystallographica Section D: Biological Crystallography 2010, 66, 12.

