## Supporting Information for "EnZight: A Structure-Guided Algorithm to Identify and Prioritize Substitution Hotspots for Enzyme Engineering"

### Experimental Procedures and Program Workflow

#### Overview of EnZight

EnZight is a Python based application for structure-guided homology analysis and hotspot determination. The full source code is available at https://github.com/morth-lab/EnZight, and EnZight can also be accessed via a web server (https://services.healthtech.dtu.dk/services/EnZight-1.0). EnZight compares a query structure (PDB or CIF format) against structural homologs, generates both a structure refined multiple sequence alignment (MSA) and a structural alignment colored by per residue conservation, and facilitates visualization in PyMOL[29].

Upon upload of a query structure, ideally with a molecular mass of at least 30 kDa to provide a sufficiently large protein core, EnZight executes four sequential modules: (1) homology search; (2) structural superposition and conservation scoring; (3) structure-refined MSA generation; and (4) hotspot determination.

##### Homology search

The query input structure is loaded by the user, after which EnZight retrieves structural homologs, by default using Foldseek[28] against the AlphaFold UniProt50 v6 database[15,16,47]. Users may change the database being searched or provide custom homologs. Hits are filtered by a user defined TM score or 3Diaa cutoff (default: TM score with a cutoff of 0.7), or by retaining the top N matches (default: 20) according to TM/3Diaa score. For optimal hotspot prediction, homologs should ideally share at least 50 % sequence identity with the query.

##### Structural superposition and residue conservation scoring

Selected homologs are superimposed onto the query using PyMOL's built in SUPER algorithm[29]. Alignments with backbone RMSD $> 5$ Å, or beyond a user specified cutoff, are discarded to retain only sufficiently similar structures for reliable residue-level comparisons.

Per-residue conservation is calculated for all structures included in the structural alignment, comprising the uploaded query structure and its selected homologs. For a given reference residue, the corresponding residue in each other structure is identified based on spatial proximity. Specifically, the residue from each other structure with the shortest C$\alpha$–C$\alpha$ distance to the reference residue is selected. If no residue is found within a predefined distance cutoff (default: 5 Å), the position is classified as a gap.

Conservation between each pair of reference and homolog residues is quantified using a BLOSUM substitution matrix, with BLOSUM62 used by default. To enable comparison across residue positions, each pairwise score is normalized relative to the self-substitution score of the reference residue:

$$S_{r,h} = \frac{B_{r,h} - B_{\min}}{B_{r,r} - B_{\min}}, \tag{S1}$$

where $S_{r,h}$ is the normalized score between reference residue $r$ and its structurally corresponding residue in homolog $h$, $B_{r,h}$ is the corresponding raw BLOSUM score, $B_{r,r}$ is the self-substitution score of reference residue $r$, and $B_{\min}$ is the minimum value in the substitution matrix. Gaps in the homolog sequence are assigned an $S_{r,h}$ value of zero.

For each reference residue, one normalized score is calculated for each of the $n$ homologs, yielding $n$ reference–homolog residue-pair scores. The conservation score for reference residue $r$ is calculated as the mean of these scores:

$$C_r = \left( \frac{1}{n} \sum_{i=1}^{n} S_{r,h} \right) \cdot 100\,\%, \tag{S2}$$

where $C_r$ is the conservation score for reference residue $r$. Conservation scores range from 0, indicating gaps or substitutions with minimal similarity across the homologs, to 100, indicating identity between the reference residue and all structurally corresponding homolog residues.

##### Structure-based multiple sequence alignment generation

An initial sequence-based multiple sequence alignment (MSA) of the query and the homologous sequences is generated using MUSCLE[48] with default parameters. The MSA is subsequently structurally refined in a two-step procedure. First, residues from different structures that align in the MSA are grouped into clusters such that all C$\alpha$ atoms within a cluster lie within a specified distance threshold (default 5 Å) of the cluster center. The cluster center is defined as the geometric center of all C$\alpha$ atoms in the cluster. If a residue falls outside this threshold, a new cluster is created. Gaps are then introduced in the MSA between clusters.

In a second step, clusters that do not align in the MSA but whose centers are closer than the same distance threshold are aligned, provided that doing so does not violate the sequential order of residues in the alignment (i.e., downstream residues cannot be aligned to upstream residues if intermediate residues are already aligned).

##### Substitution hotspot determination

Herein, substitution hotspots are defined as amino acid substitutions from the query sequence to a homologous amino acid that satisfy a set of structural and conservation-based criteria.

By default, EnZight restricts hotspot detection to the protein core. Core residues in EnZight are defined as residues belonging to the top 50 % of per-residue conservation scores, as well as residues located within 6 Å ($C\alpha$–$C\alpha$ distance) of these highly conserved positions, provided they are not surface-exposed.

For each core residue in the query structure ($r_q$) and its corresponding residue in a homologous structure ($r_h$), the local structural cavity is evaluated by mapping neighboring residues. A neighboring residue is defined as a residue in the query whose sidechain–to–sidechain distance to $r_q$ or $r_h$ is $\leq 4$ Å. A position is considered a substitution hotspot if all of the following conditions are met: (i) the amino acid identity of the query residue differs from that of the homolog residue, (ii) all mapped neighboring residues in the homolog are identical to, or larger than, their query counterparts according to predefined substitution rules (G→all; A→all except G/P; S→C/T; V→I; F→Y), and (iii) at least one neighbor residue was found.

When these criteria are satisfied, the substitution of the query residue to the corresponding homolog amino acid is designated as a substitution hotspot. EnZight additionally supports the identification of double-mutant hotspots by detecting pairs of residues within the same local structural environment that combined satisfy the hotspot criteria.

Hotspots are ranked according to the EnZight score, defined as

$$E_{\text{score}} = (100 - \text{mean}\,(C_{\text{sub}})) + (100 - \text{mean}\,(C_{\text{neighbor}}))\,, \quad \text{(S3)}$$

where $E_{\text{score}}$ is the EnZight score, $C_{\text{sub}}$ denotes the conservation score of the position(s) subjected to substitution in the query structure, and $C_{\text{neighbor}}$ denotes the conservation scores of residues in the local structural environment surrounding the target position or positions. Thus, higher EnZight scores indicate lower conservation of both the substituted position(s) and their local structural environment.

#### Experimental validation

The bacterial urethanase u10 (NCBI ID: WP_265550772.1) from *Sphingomonas* sp. S1-29 was used as query to a NCBI blastp search[49,50]. It was searched against all databases available on NCBI blast by August 2025. Sequence identity of 60 % to the query was used as a cut off. AlphaFold 3 structures were made of the query sequence and all the entries above the cut off using the AlphaFold 3 server[15]. Predicted structures were used as query and homologs in EnZight to identify substitution hotspots in u10.

##### Plasmid variant library construction

Gene encoding u10 was codon optimized for expression in *Escherichia coli* (Genscript, Table S3) and commercially synthesized in a pETM11 backbone (Genscript). The gene was incorporated into the plasmid using the NcoI and XhoI restriction sites introducing a His-tag and a TEV cleavage site in the N-terminal of the protein. pETM11 contains a gene coding for kanamycin resistance and, therefore, kanamycin ($50\,\mu g\,mL^{-1}$) was used for selection of cells containing pETM11.

Variants of u10 were constructed by amplifying and linearizing full length plasmid by PCR using CloneAmp HiFi PCR Premix (TAKARA) with partially overlapping primers (Table S4). Primers were designed using the principle of the In-Fusion Cloning System primer design tool (TAKARA). FastDigest DpnI (Thermo Scientific) treated PCR products were purified with GeneJET PCR Purification Kit (Thermo Scientific) and transformed into *E. coli* DH5$\alpha$ (Invitrogen) using the Mix & Go *E. coli* Transformation Kit (Zymo research). Plasmids were purified from overnight cultures of the transformed cells using GeneJET Plasmid Miniprep Kit (Thermo Scientific), verified by Sanger sequencing (Eurofins Genomics), and stored at −20 °C.

##### Protein expression and purification

Plasmids from a variant library were transformed into *E. coli* BL21 (DE3) (Invitrogen) using the Mix & Go *E. coli* Transformation Kit (Zymo research). From a preculture of the transformed cells in LB, a 1/100 dilution were made in fresh LB (50 mL for small scall and 1 L for large scale) and incubated at 37 °C in a shaking flask with plenty aeration. At $OD_{600}$ at approximately 0.4–0.6, a final concentration of 100 µM Isopropyl $\beta$-D-1-thiogalactopyranoside (IPTG) was added to the culture and the culture was incubated at 15 °C with shaking for 24 hours. The cells were harvested by centrifugation (4000 × g, 15 min, 4 °C) and stored at −20 °C.

For cell lysis, frozen cell pellets were resuspended in ice cold lysis buffer (300 mM NaCl, 50 mM Tris-HCl, pH 8.0) in a weight ratio of approximately 1:10 (pellet:buffer). The solution was supplemented with a final concentration of 2 µg $mL^{-1}$ bovine pancreas DNase I (Merck) and a final concentration of 0.1 mg $mL^{-1}$ Lysozyme from chicken egg white (Merck). Pellets were completely resuspended using vortex. For small scale expressions, cells were lysed by sonication using a Q500 Sonicator (Qsonica) at 35 % amplitude. The total sonication time was 1 minute ON with cycles of 5 sec ON and 9 sec OFF. Cell debris was removed using centrifugation (15 000 × g, 15 min, 4 °C). For large scale expression, cells were lysed by homogenization using a Pressure Cell Homogeniser from Stansted (1 bar pressure) and cell debris was removed using centrifugation (30 000 × g, 30 min, 4 °C).

Recombinant proteins were purified from lysates by nickel affinity chromatography. Cell lysates were applied to equilibrated Ni–NTA resin either by gravity flow or using an automated chromatography system, depending on expression scale. For gravity-flow purification, lysates were gently loaded onto PD-10 columns (Cytiva) packed with 1.5–2 mL HisPur™ Ni-NTA resin (Thero Fisher Scientific). Columns were washed with five column volumes (CV) of wash buffer 1 (10 mM imidazole, 500 mM NaCl, 50 mM Tris-HCl, pH 8.0), followed by two CV of wash buffer 2 (10 mM imidazole, 50 mM Tris-HCl, pH 8.0). Proteins were eluted with five CV of elution buffer (300 mM imidazole, 300 mM NaCl, 50 mM Tris-HCl, pH 8.0). For larger-scale purifications, lysates were loaded onto a 5 L HisTrap Ni-NTA column (Cytiva) using an ÄKTA chromatography system (Cytiva). After washing with wash buffers 1 and 2, proteins were eluted using a linear gradient of elution buffer. Elution fractions containing the target protein were pooled.

Eluted proteins were upconcentrated and the buffer was changed to storage buffer (150 mM NaCl, 10 mM Tris-HCl, pH 8.0) using Amicon Ultra centrifugal filters (30 kDa MWCO, Millipore) according to the manufacturer's instructions. Proteins for crystallization were additionally purified by size-exclusion chromatography on a HiLoad Superdex 200 pg 10/600 column (Cytiva). Proteins were snap-frozen in liquid nitrogen and stored at −80 °C.

##### Protein stability and activity assays

Protein stability was assayed by measuring apparent protein unfolding temperature ($T_{m,app}$). 0.5 mg $mL^{-1}$ purified protein was analyzed using the Prometheus Panta equipment (Nanotemper). The temperature was increased 1.5 °C per minute from 20 °C to 90 °C. $T_{m,app}$ was estimated as the temperature where the first derivative of 330 nm was at its minimum. This was determined using the Panta analysis software (Nanotemper).

Enzyme activity was assayed by measuring the conversion of di-urethane ethylene methylenedianiline (DUE-MDA), flutamide, or four *para*-nitrophenyl ($p$NP) substrates: $p$NP-butyrate, $p$NP-butanamide, ethyl $p$NP-carbamate, and 1-ethyl-3-$p$NP-urea. DUE-MDA and its hydrolysis product, mono-urethane ethylene methylenedianiline (MUE-MDA), were synthesized as described previously [51].

Reaction mixtures (200 µL) contained 30–5000 nM enzyme and 0.016–1.14 mM substrate in 10 mM Tris-HCl and 150 mM NaCl at pH 8.0. Substrate stock solutions were prepared in 96 % ethanol, resulting in a final ethanol concentration of 4.8 % (v/v) in the reaction mixture. DUE-MDA assays were performed in a benchtop thermomixer, whereas the remaining assays were performed in a plate reader. All assays were conducted at 40 °C with shaking.

The enzymatic conversion of flutamide and the $p$NP substrates was monitored colorimetrically by measuring absorbance at 405 nm at 1-min intervals. Product formation was quantified using the corresponding product standard curves (Figure S2a). Conversion of DUE-MDA to MUE-MDA was quantified by HPLC as described previously [31] using an MUE-MDA standard curve (Figure S2b).

Initial rates were determined from the linear regions of the progress curves. DUE-MDA assays used for Michaelis–Menten analysis were performed over the indicated substrate concentration range, whereas endpoint assays were performed using 1 mM DUE-MDA. Each condition was assayed in technical triplicate. Data processing and quantification were performed using Chromeleon, Microsoft Excel, and Python. The kinetic parameters $k_{cat}$ and $K_m$ were determined by nonlinear regression using the `curve_fit` function from the SciPy package in Python. Correlations between EnZight scores and activity enhancement were evaluated using two-sided Spearman rank correlation in SciPy, with statistical significance defined as $p < 0.05$.

##### Crystallization and structure determination

The triple variant u10_ITP was incubated with the suicide inhibitor LUI [37,38] at a 1:1 molar ratio for 1 hour with inversion. LUI (Figure S4) was synthesized as described previously [38]. The u10_ITP:LUI complex and u10_WT were crystallized with sitting drop vapor diffusion with a concentration of 22 mg $mL^{-1}$ and 17 mg $mL^{-1}$, respectively. Mosquito Xtal3 (SPT Labtech) robot was used to set up crystal plates. Initial screening was conducted with different commercial kits in SwissCi XTAL (SD-3) 3 well plates, yielding a single crystal of u10_ITP in the condition with 10 % (v/v) 2-propanol, 25 % (w/v) PEG smear broad, and 0.1 M Bicine (pH 9.0), and a single crystal of u10_WT in the condition with 0.2 M Sodium Malonate and 12 % (w/v) PEG 3350 (pH 7.0). Optimization of the u10_ITP crystal condition was done in MRC Maxi 48 Well plates. The optimal condition for yielding large plate-like crystals of u10_ITP at room temperature within 96 hours was 10 % (v/v) 2-propanol, 20 % (w/v) PEG smear broad, and 0.1 M Bicine (pH 9.0). All crystals were found in wells with 1:1 (volumetric ratio) protein:reservoir drops. 15 % (v/v) glycerol was used for cryoprotection before flash-freezing in liquid $N_2$, and X-ray diffraction data collected. Optimization of the u10_WT crystal condition was done in 24-well plates with different ratios of seed stocks varying from 1:10 to 1:1000 of undiluted seed stock: mother liquor. 20 % (v/v) glycerol was used for cryoprotection before flash-freezing in liquid $N_2$, and X-ray diffraction data collected. The data reduction, space group determination and merging were done by XDS, pointless and XSCALE respectively [52,53]. The molecular replacement solutions were found with Phaser using an AlphaFold 3 prediction of u10 as a search model [15,54]. From this, the crystal structure was built and refined using Coot and phenix.refine [55,56]. Ramachandran plots and further validation were performed with MolProbity [57], and figures were constructed using PyMOL [29]. All models and structure factors are deposited to the Protein Data Bank (PDB).

**Table S1.** Experimental characterization of u10 EnZight variants. DUE–MDA end-point activity is reported as the measured concentration of MUE–MDA. Flutamide activity is reported as the initial velocity normalized to enzyme concentration. Values are reported as mean ± standard deviation.

| Variant | EnZight score | Cloned | Expressed | DUE–MDA activity [MUE–MDA] (mM) | Flutamide activity $V_0/E_0$ ($s^{-1}$) | $T_{m,app}$ (°C) | $n$[a] |
|---|---|---|---|---|---|---|---|
| WT | – | Yes | Yes | $0.25 \pm 0.05$ | $1.49 \pm 0.06$ | $59.69 \pm 0.19$ | 3 |
| G158S | 65.761 | Yes | Yes | $0.27 \pm 0.04$ | $1.68 \pm 0.07$ | $58.50 \pm 0.05$ | 1 |
| C399V | 63.462 | Yes | Yes | $0.31 \pm 0.05$ | $2.71 \pm 1.10$ | $59.56 \pm 0.01$ | 1 |
| C399T | 63.462 | Yes | Yes | $0.31 \pm 0.05$ | $3.30 \pm 0.23$ | $53.85 \pm 0.01$ | 1 |
| S151P | 62.500 | Yes | Yes | $0.44 \pm 0.02$ | $2.46 \pm 0.24$ | $59.95 \pm 0.24$ | 3 |
| S151C | 62.500 | Yes | Yes | $0.25 \pm 0.02$ | $0.21 \pm 0.03$ | $60.51 \pm 0.11$ | 3 |
| H90T | 54.163 | Yes | Yes | $0.32 \pm 0.02$ | $1.50 \pm 1.22$ | $60.54 \pm 0.12$ | 3 |
| A27C | 53.043 | Yes | Yes | $0.22 \pm 0.07$ | $1.25 \pm 0.21$ | $59.89 \pm 0.02$ | 1 |
| A27T | 53.043 | Yes | Yes | $0.24 \pm 0.03$ | $2.11 \pm 0.36$ | $59.94 \pm 0.18$ | 1 |
| M400V | 49.396 | Yes | Yes | $0.27 \pm 0.02$ | $2.73 \pm 1.75$ | $59.15 \pm 0.01$ | 1 |
| M400F | 49.396 | Yes | Yes | $0.16 \pm 0.04$ | $0.95 \pm 0.15$ | $59.52 \pm 0.01$ | 1 |
| T416C | 40.550 | Yes | Yes | $0.18 \pm 0.03$ | $2.34 \pm 0.27$ | $60.65 \pm 0.12$ | 1 |
| S122T | 34.420 | Yes | Yes | $0.10 \pm 0.02$ | $0.35 \pm 0.02$ | $61.17 \pm 0.02$ | 1 |
| I232T | 33.722 | Yes | Yes | $0.10 \pm 0.00$ | $0.50 \pm 0.04$ | $57.43 \pm 0.01$ | 1 |
| I232V | 33.722 | Yes | Yes | $0.20 \pm 0.03$ | $1.13 \pm 0.56$ | $58.50 \pm 0.03$ | 1 |
| V265C | 32.246 | Yes | Yes | $0.10 \pm 0.02$ | $1.17 \pm 0.16$ | $57.69 \pm 0.02$ | 1 |
| M231L | 25.523 | Yes | No | – | – | – | – |
| L221M | 22.935 | Yes | Yes | $0.15 \pm 0.01$ | $0.56 \pm 0.16$ | $59.93 \pm 0.00$ | 1 |
| I415M | 21.264 | Yes | Yes | $0.17 \pm 0.05$ | $1.85 \pm 0.22$ | $60.33 \pm 0.00$ | 1 |
| I415L | 21.264 | Yes | Yes | $0.25 \pm 0.03$ | $0.82 \pm 0.06$ | $58.55 \pm 0.00$ | 1 |
| L171M | 11.451 | Yes | Yes | $0.13 \pm 0.02$ | $1.15 \pm 0.63$ | $55.92 \pm 0.02$ | 1 |
| A397S | 11.413 | Yes | Yes | $0.16 \pm 0.06$ | $2.15 \pm 0.53$ | $58.85 \pm 0.03$ | 1 |
| A397V | 11.413 | No | No | – | – | – | – |
| I34V | 11.329 | Yes | Yes | $0.11 \pm 0.01$ | $0.52 \pm 0.02$ | $57.25 \pm 0.02$ | 1 |
| V44I | 10.618 | Yes | Yes | $0.24 \pm 0.01$ | $0.74 \pm 0.25$ | $60.74 \pm 0.30$ | 3 |
| I218V | 9.330 | Yes | Yes | $0.19 \pm 0.00$ | $1.02 \pm 0.07$ | $60.24 \pm 0.03$ | 1 |
| I169L | 7.419 | Yes | Yes | $0.25 \pm 0.08$ | $2.19 \pm 0.05$ | $58.93 \pm 0.01$ | 1 |
| V124I | 6.067 | Yes | Yes | $0.16 \pm 0.02$ | $0.71 \pm 0.13$ | $59.94 \pm 0.04$ | 1 |
| V124L | 6.067 | Yes | Yes | $0.02 \pm 0.00$ | $0.02 \pm 0.00$ | $59.66 \pm 0.01$ | 1 |
| V349M | 5.761 | Yes | Yes | $0.20 \pm 0.07$ | $1.13 \pm 0.19$ | $57.89 \pm 0.06$ | 1 |
| L389I | 5.512 | Yes | Yes | $0.21 \pm 0.05$ | $1.83 \pm 0.03$ | $59.37 \pm 0.09$ | 1 |
| L389V | 5.512 | Yes | Yes | $0.14 \pm 0.03$ | $1.71 \pm 0.29$ | $60.33 \pm 0.00$ | 1 |
| M76I | 4.734 | Yes | Yes | $0.04 \pm 0.00$ | $0.09 \pm 0.03$ | $55.29 \pm 0.04$ | 1 |
| P125A | 4.521 | Yes | Yes | $0.11 \pm 0.04$ | $0.35 \pm 0.02$ | $58.60 \pm 0.04$ | 1 |
| I117V | 4.348 | Yes | Yes | $0.24 \pm 0.03$ | $1.51 \pm 0.11$ | $59.65 \pm 0.01$ | 1 |
| L371I | 3.221 | No | No | – | – | – | – |
| V181I | 3.043 | Yes | Yes | $0.12 \pm 0.01$ | $0.90 \pm 0.16$ | $60.64 \pm 0.06$ | 1 |
| V44I_H90T | – | Yes | Yes | $0.28 \pm 0.07$ | $3.69 \pm 0.42$ | $61.47 \pm 0.09$ | 2 |
| V44I_S151P | – | Yes | Yes | $0.43 \pm 0.01$ | $3.37 \pm 0.46$ | $61.13 \pm 0.17$ | 2 |
| V44I_S151C | – | Yes | Yes | $0.17 \pm 0.01$ | $1.19 \pm 0.16$ | $61.65 \pm 0.06$ | 2 |
| H90T_S151P | – | Yes | Yes | $0.47 \pm 0.01$ | $3.72 \pm 0.37$ | $60.92 \pm 0.06$ | 2 |
| H90T_S151C | – | Yes | Yes | $0.28 \pm 0.02$ | $1.55 \pm 0.35$ | $61.42 \pm 0.03$ | 2 |
| V44I_H90T_S151P | – | Yes | Yes | $0.47 \pm 0.02$ | $3.46 \pm 0.66$ | $61.77 \pm 0.09$ | 2 |
| V44I_H90T_S151C | – | Yes | Yes | $0.28 \pm 0.02$ | $1.89 \pm 0.52$ | $62.15 \pm 0.13$ | 2 |

[a] Number of independently constructed and characterized variants.

**Table S2.** Data collection and refinement statistics for the u10 wild-type and u10 ITP structures. Values in parentheses refer to the highest-resolution shell.

| Structure | u10 wild-type | u10 ITP |
|---|---|---|
| PDB ID | 32SM | 32SK |
| Beamline | BioMAX | BioMAX |
| Wavelength (Å) | 0.7293 | 0.7293 |
| Resolution (Å) | 23.72–1.81 (1.83–1.81) | 48.71–2.24 (2.32–2.24) |
| Space group | $P\,2_1 2_1 2_1$ | $P\,6_3 22$ |
| $a$, $b$, $c$ (Å) | 72.605, 104.643, 119.113 | 105.165, 105.165, 172.942 |
| $\alpha$, $\beta$, $\gamma$ (°) | 90, 90, 90 | 90, 90, 120 |
| Total reflections | 1 060 120 (21 244) | 351 329 (17 728) |
| Unique reflections | 82 995 (2 651) | 27 807 (2 642) |
| $R_{\mathrm{merge}}$ | 0.1171 (> 1) | 0.2812 (> 1) |
| $R_{\mathrm{meas}}$ | 0.122 (> 1) | 0.2928 (> 1) |
| $R_{\mathrm{pim}}$ | 0.0339 (> 1) | 0.0792 (0.425) |
| Multiplicity | 12.8 (8.0) | 12.6 (6.7) |
| Completeness (%) | 98.49 (87.95) | 99.63 (97.52) |
| $I/\sigma(I)$ | 12.83 (0.45) | 7.46 (1.64) |
| Wilson $B$-factor (Å$^2$) | 40.40 | 27.33 |
| $\mathrm{CC}_{1/2}$ | 0.996 (0.0677) | 0.975 (0.534) |
| | Refinement statistics | |
| Reflections used in refinement | 82 000 (2 482) | 27 788 (2 640) |
| Reflections used for $R_{\mathrm{free}}$ | 4 130 (139) | 1 438 (160) |
| $R_{\mathrm{work}}$ | 0.2063 (0.5086) | 0.1791 (0.2527) |
| $R_{\mathrm{free}}$ | 0.2358 (0.5076) | 0.2138 (0.2667) |
| No. of non-hydrogen atoms | 5 697 | 3 489 |
| Macromolecules | 5 474 | 3 177 |
| Ligands | 0 | 27 |
| Solvent | 223 | 285 |
| Protein residues | 750 | 432 |
| RMSD, bond lengths (Å) | 0.010 | 0.050 |
| RMSD, bond angles (°) | 1.08 | 0.62 |
| Ramachandran favored (%) | 96.19 | 97.21 |
| Ramachandran allowed (%) | 3.41 | 2.79 |
| Ramachandran outliers (%) | 0.40 | 0.00 |
| Rotamer outliers (%) | 1.29 | 0.32 |
| Clashscore | 5.46 | 1.41 |
| Average $B$-factor (Å$^2$) | 65.62 | 31.01 |
| Macromolecules | 65.91 | 30.63 |
| Ligands | – | 52.75 |
| Solvent | 58.48 | 33.13 |

**Table S3.** Gene sequence used in this study.

| Gene name | DNA sequence |
|---|---|
| u10<br>NCBI ID:<br>WP_265550772.1 | atgaaacatcaccatcaccatcaccccatgagcgattacgacatccccactactgagaatcttta<br>ttttcagggcgccatggcgacttcaacaccacccctaacggctcacgcaacggctgctgctatcc<br>gcgcgggcacgaccaccgcaagagaacaggcattggcggcgatcgcgcgcattgagacgctggat<br>ccggagatcaacgccgtcccggtgcgtgactttgatcgcgcactagcagccgctgacgcagcgga<br>tgcccgtctggctgcgggcgacacagcgcctttgctgggtgttccgatgaccgtgaaagaagcat<br>tcgatatcgcagggctgccaacgcattggggttttgcggaacatcgtaacaacgtggcgaccacc<br>gacgctcacgccgttgcaaggctgaaggccgcaggcgcgattatcctgggtaaaagcaacgtgcc<br>gaaaggtctgggtgattggcagagcgttaacagcattcacggtcgtacgaaccacccgttggacc<br>cggctcgtacttcgggtggctctagcgggggcggagcagcggcggttgccgcgggcatggtgccg<br>attgagctgggctcggacgtgggtggctccattcgtgtcccggcacattttgtggtatctgggg<br>tcataagccgagctggaatgcaatcagcagcgatggtcatcgttatccgggcactgacggtactg<br>agaccgttctgggcgtcattggtccgctggcgcgcgacccgcaggatctggcgctgatgattgat<br>ctgttggcgaccctgccgctgccgcgtccggccgctcgcacccgccgtgtactggttctggccca<br>acatccagagacccgtaccgcccatgcagtggtggaaggtgttgaacgtgcggcggctgccctag<br>ccagagccggcgttgaggttgttcgtcattctgaccttctcccagatctgagccgtcagcacagc<br>gcatatggtgacttgctgaatgtgaccttcgcgcgctccaatccggcgcttcaccacaccctccc<br>gagcctgctcaagtggctgagtatgctggacgcgcaagcgcgttccacccgcgcttggggcgcct<br>tgtttggtgaattcgatgccgtaatcgcgccaccggcggctacccaggcgttcgcgcacgatcac<br>agcccgttggcggaccgtaccttagcgattgacggcaccccgggtccgtacgatgcacacttggc<br>gtgggcaggccttgcgacctacccgggtctgccagcgacgtgcatgccggttggtttaatcgacg<br>gcctgccgaccggtgtgcaggtcatcaccgaactgcaccaagatcatcgtgctatcgagattgcg<br>gcactgatcgcacaacacctgtccccgtctccgcaaggtgccatcgcgtaa |

**Table S4.** Primers used for site-directed mutagenesis. Primer sequences are given in the $5'$ to $3'$ direction. For each target exchange, the reverse primer is listed first, followed by the forward primer.

| **Target exchange** | **Sequence ($5'$ to $3'$), reverse/forward** |
|---|---|
| S178A[a] | gaatggcgccacccacgtccgagc<br>tgggtggcgccattcgtgtcccggcacat |
| T12I | ttgcgtgagccgttaggggt<br>taacggctcacgcaatcgctgctgctatccgcgc |
| A27C | ctgttctcttgcggtggtcgtg<br>accgcaagagaacagtgcttggcggcgatcgc |
| A27T | ctgttctcttgcggtggtcgtg<br>accgcaagagaacagacattggcggcgatcgc |
| I34V | gatccagcgtctcaacacgtgcaatcgccgccaatgcctg<br>ttgagacgctggatccggagatc |
| V44I | ggcgttgatctccggatccag<br>ccggagatcaacgccatcccggtgcgtgactttgatc |
| M76I | atcggaacacccagcaaaggc<br>gctgggtgttccgattaccgtgaaagaagcattcgatatcg |
| H90S | ttccgcaaaaccccagctcgttggcagccctgcgatat<br>tggggttttgcggaacatcgtaac |
| H90T | ttccgcaaaaccccaggtcgttggcagccctgcgatat<br>tggggttttgcggaacatcgtaac |
| I117V | ttttacccaggataactgcgcctgcggccttcag<br>ttatcctgggtaaaagcaacgtgcc |
| S122T | cacccagacctttcggaacattcgttttacccaggataatcgcgcctg<br>cgaaaggtctgggtgattggcag |
| V124L | cacccagacctttcggcaggttgcttttacccaggataatcg<br>cgaaaggtctgggtgattggcag |
| V124I | cacccagacctttcggaatgttgcttttacccaggataatcg<br>cgaaaggtctgggtgattggcag |
| P125A | cacccagacctttcgccacgttgcttttacccaggataatcg<br>cgaaaggtctgggtgattggcag |

Continued on the next page

Table S4 continued from the previous page

| Target exchange | Sequence (5′ to 3′), reverse/forward |
|---|---|
| H143N | gttcgtacgaccgtgaatgctgttaac<br>cacggtcgtacgaacaacccattggacccggctcgtactt |
| S151C | agtacgagccgggtccaacg<br>gacccggctcgtacttgtggtggctctagcggg |
| S151P | agtacgagccgggtccaacg<br>gacccggctcgtactccaggtggctctagcggg |
| V162L | atgcccgcggccagcgccgctgctccgcc<br>gctggccgcgggcatggtgc |
| I169L | ccagctccagcggcaccatgcccgcggcaa<br>tgccgctggagctgggctcggacgtggg |
| L171M | ccatctcaatcggcaccatgccc<br>tgccgattgagatgggctcggacgtgggtg |
| V181I | aaaatgtgccgggatacgaatggagccacccacg<br>atcccggcacattttgtggtatctg |
| I218V | ccagcggacccacgacgcccagaacggtctcag<br>tcgtgggtccgctggcgcgc |
| L221M | ccatcggaccaatgacgcccagaac<br>tcattggtccgatggcgcgcgacccg |
| M231L | cagcgccagatcctgcgg<br>caggatctggcgctgttaattgatctgttggcgaccctgc |
| M231V | cagcgccagatcctgcgg<br>caggatctggcgctggtgattgatctgttggcgaccctg |
| I232V | cagcgccagatcctgcgg<br>caggatctggcgctgatggttgatctgttggcgaccctgc |
| I232T | cagcgccagatcctgcgg<br>caggatctggcgctgatgaccgatctgttggcgaccctgc |
| I232L | cagcgccagatcctgcgg<br>caggatctggcgctgatgctggatctgttggcgaccctgc |
| V250A | gccagaaccagtgcacggcgggtgcgagc<br>tgcactggttctggcccaacatccag |
| V265C | cttccacacatgcatgggcggtacgggt<br>atgcatgtgtggaaggtgttgaacgtgcg |
| V349M | ggcatcgaattcaccaaacaagg<br>ggtgaattcgatgccatgatcgcgccaccggcggcta |
| I350L | ggcatcgaattcaccaaacaagg<br>ggtgaattcgatgccgtactggcgccaccggcggcta |
| L389I | ccgggtaggtcgcaatgcctgcccacgccaagtg<br>ttgcgacctacccgggtctg |
| L389V | ccgggtaggtcgcaacgcctgcccacgccaagtg<br>ttgcgacctacccgggtctg |
| A397S | gattaaaccaaccggcatgcacgtgcttggcagacccgggtaggtc<br>ccggttggtttaatcgacggcct |
| C399V | gattaaaccaaccggcatgaccgtcgctggcagacccg<br>ccggttggtttaatcgacggcct |
| C399T | gattaaaccaaccggcatggtcgtcgctggcagacccg<br>ccggttggtttaatcgacggcct |
| M400V | gattaaaccaaccggcacgcacgtcgctggcagacc<br>ccggttggtttaatcgacggcct |
| M400F | gattaaaccaaccggaaagcacgtcgctggcagacccg<br>ccggttggtttaatcgacggcct |
| T410G | atgacctgcacaccgcccggcaggccgtcgattaaacc<br>cggtgtgcaggtcatcaccgaa |
| T410A | atgacctgcacaccggccggcaggccgtcgattaaacc<br>cggtgtgcaggtcatcaccgaa |
| V412A | atgacctgcgcaccggtcggcaggcc<br>cggtgcgcaggtcatcaccgaactgcac |

Continued on the next page

Table S4 continued from the previous page

| Target exchange | Sequence (5′ to 3′), reverse/forward |
|---|---|
| I415L | ttcggtcaggacctgcacaccggtcgg<br>caggtcctgaccgaactgcaccaagatcatcgtg |
| I415M[b] | agttcggUcatgacctgcacaccggtcg<br>accgaacUgcaccaagatcatcgtg |
| T416C | ttcgcagatgacctgcacaccggtcg<br>caggtcatctgcgaactgcaccaagatcatcgtgc |
| T416A | ttcggcgatgacctgcacaccggtcg<br>caggtcatcgccgaactgcaccaagatcatcgtgc |
| A424S | cgatagaacgatgatcttggtgcagttcggtg<br>atcatcgttctatcgagattgcggcactgatcg |
| I427A | gccgcagcctcgatagcacgatgatcttggtgcag<br>tatcgaggctgcggcactgatcgcacaac |

[a] Alanine mutation of catalytic serine for inactive variant.
[b] Primers for USER-cloning.

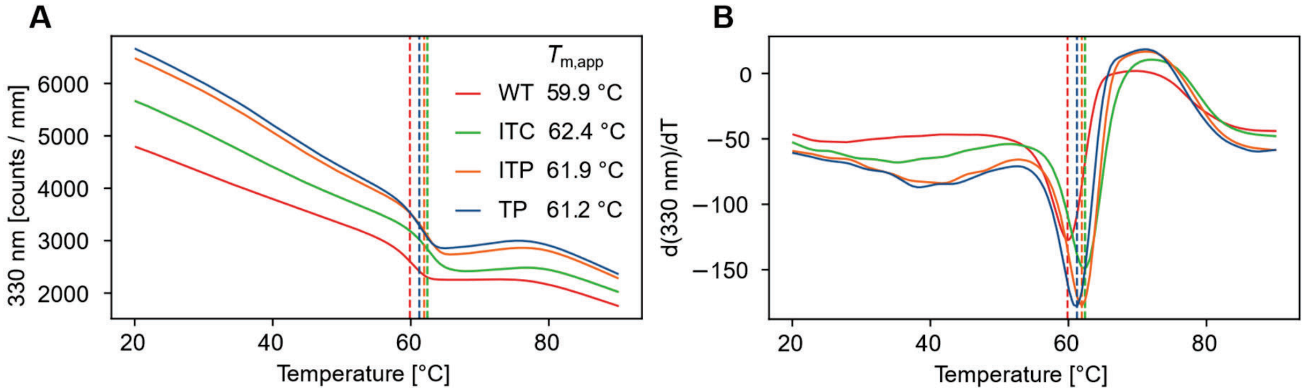


**Figure S1.** NanoDSF thermal unfolding data for u10 wild-type and selected variants. Vertical dashed lines show inflection points which are also noted as $T_{\text{m,app}}$. a) 330 nm fluorescence. b) First derivative of 330 nm fluorescence.

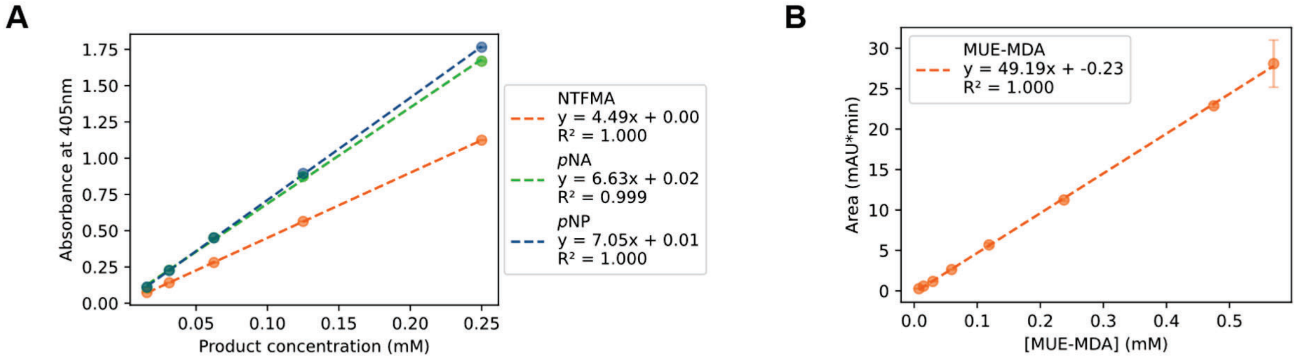


**Figure S2.** Standard curves. a) Colorimetric product standard of 4-Nitro-3-(trifluoromethyl)aniline (NTFMA), *para*-Nitroaniline (*p*NA), and *para*-Nitrophenol (*p*NP). b) HPLC product standard mono-urethane ethylene methylenedianiline (MUE-MDA). All data points are reported as mean ± SD from technical triplicates.

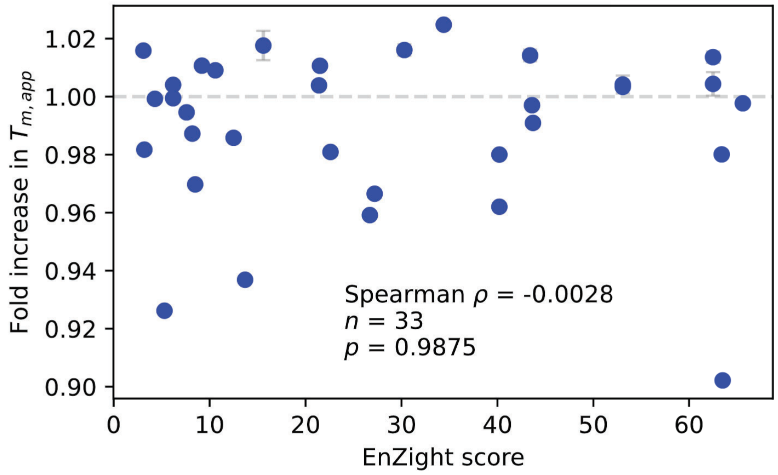


**Figure S3.** Correlation between $T_{\mathrm{m,app}}$ and EnZight score for all tested u10 variants. All data points are reported as mean ± SD from technical triplicates. Statistical parameters (Spearman $\rho$, number of samples (n), p-value) for the correlation were included. Horizontal line ($y = 1$) was included as wild-type activity.

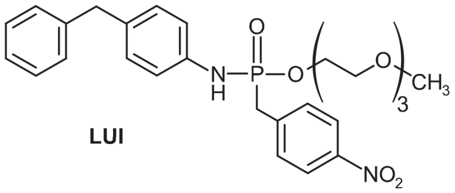


**Figure S4.** Chemical structure of the suicide inhibitor LUI.

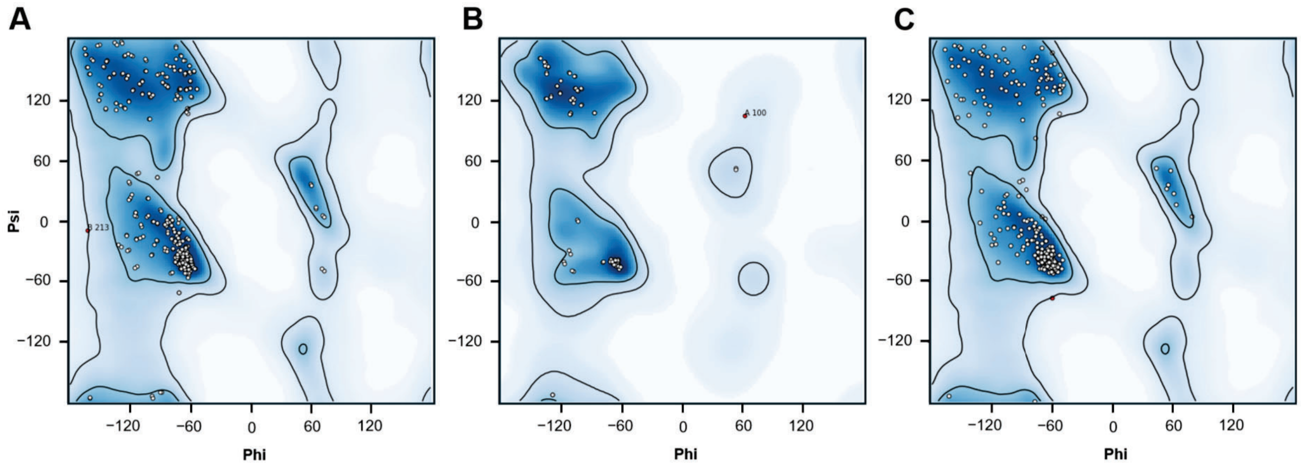


**Figure S5.** Ramachandran plots showing outliers. a) Non-pro/gly residues in u10 wild-type. Residue 213, chain B shown in red. b) Ile/leu residues u10. Residue 100, chain A shown in red. c) Non-pro/gly residues in u10 ITP variant with residue 244 shown in red.

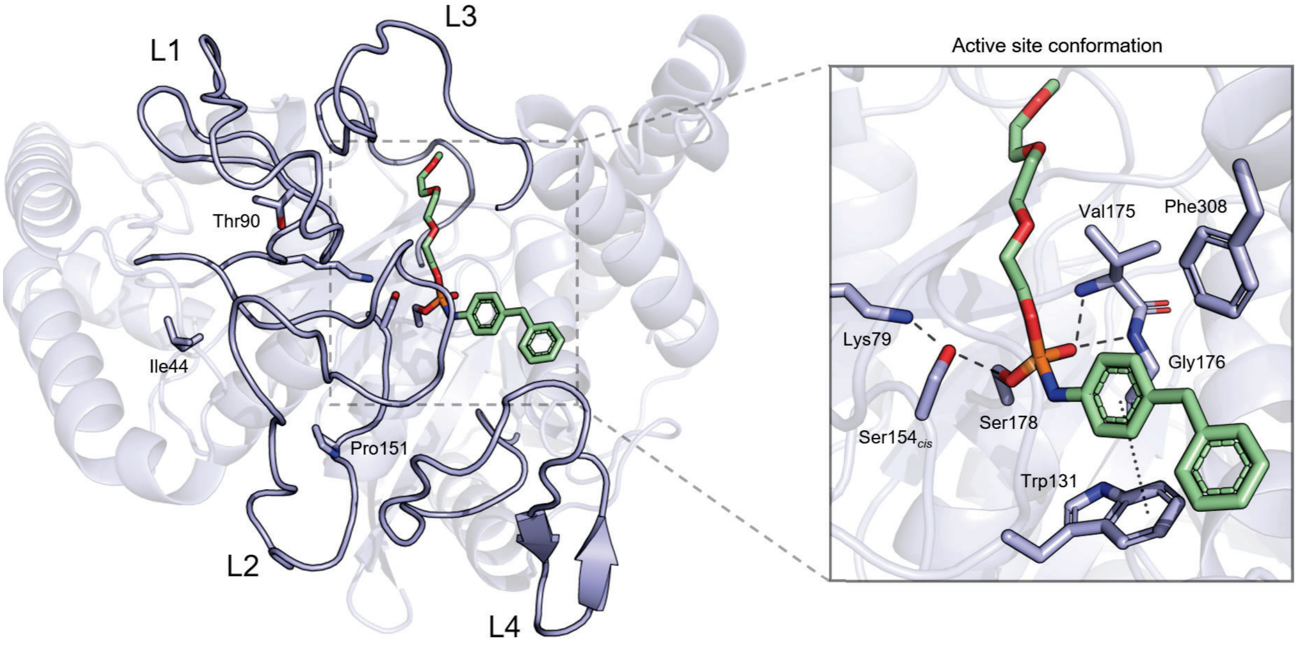


**Figure S6.** Structural overview of the u10 ITP variant in complex with the amidase signature inhibitor LUI (green). Amidase signature loops highlighted (L1-L4). The catalytic triad and mutated residues are displayed as sticks. The magnified view shows the active site with key residues as sticks. $\pi$-$\pi$ interaction is indicated by a dotted line ($4.5\,\text{Å}$) and hydrogen bonds are shown as dashed lines ($2.6$-$3.2\,\text{Å}$).

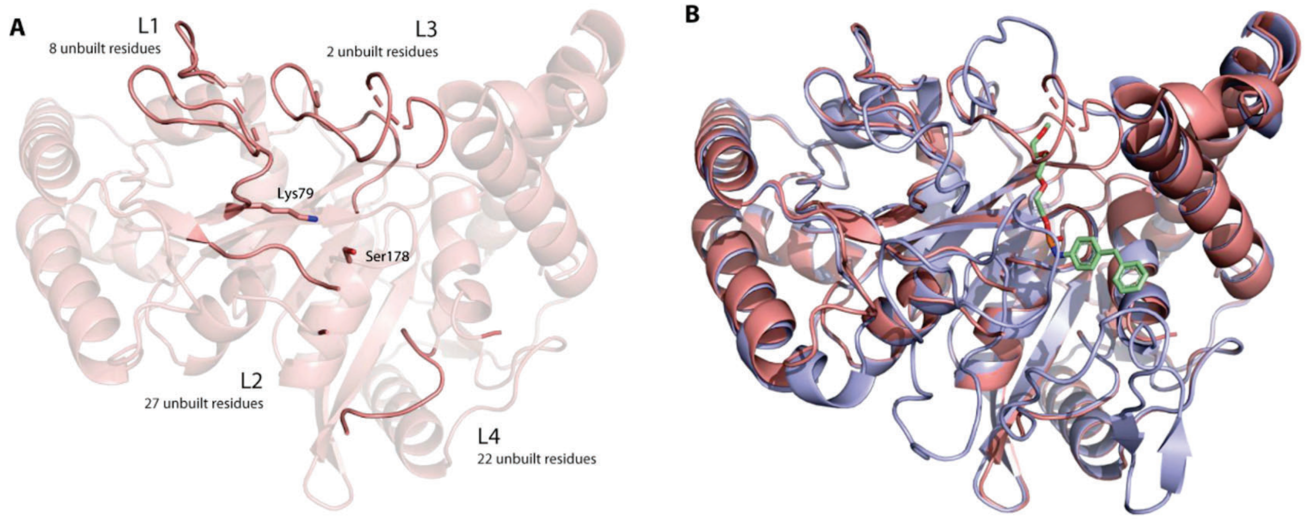


**Figure S7.** Structural overview of the u10 wild-type. a) Overall structure of the u10 wild-type (red) with amidase signature loops highlighted (L1–L4). Unbuilt residues are indicated next to corresponding loop region. Side chains of the catalytic triad residues that were built are displayed as sticks. b) Structural alignment of u10 wild-type and u10_ITP (blue) with the suicide inhibitor LUI (green).
